# Predatory fireflies and their toxic firefly prey have evolved distinct toxin resistance strategies

**DOI:** 10.1101/2023.03.08.531760

**Authors:** Lu Yang, Flora Borne, Anja Betz, Matthew L. Aardema, Ying Zhen, Julie Peng, Regina Visconti, Mariana Wu, Bartholomew P. Roland, Aaron D. Talsma, Mike J. Palladino, Georg Petschenka, Peter Andolfatto

## Abstract

Toxic cardiotonic steroids (CTS) act as a defense mechanism in many firefly species (Lampyridae) by inhibiting a crucial enzyme called Na^+^,K^+^-ATPase (NKA). While most fireflies produce these toxins internally, species of the genus *Photuris* acquire them from a surprising source: predation on other fireflies. The contrasting physiology of toxin exposure and sequestration between *Photuris* and other firefly genera suggests that distinct strategies may be required to prevent self-intoxication. Our study demonstrates that both *Photuris* and their firefly prey have evolved highly-resistant NKAs. Using an evolutionary analysis of the specific target of CTS (ATPα) in fireflies, and gene-editing in *Drosophila*, we find that the initial steps towards resistance were shared among *Photuris* and other firefly lineages. However, the *Photuris* lineage subsequently underwent multiple rounds of gene duplication and neofunctionalization, resulting in the development of ATPα paralogs that are differentially expressed and exhibit increasing resistance to CTS. In contrast, other firefly species have maintained a single copy. Our results implicate gene duplication as a facilitator in the transition of *Photuris* to its distinct ecological role as predator of toxic firefly prey.

**One-Sentence Summary:** Gene duplication and neofunctionalization distinguish firefly predators from their toxic firefly prey.

## Introduction

Many species of fireflies (family Lampyridae, subfamily Lampyrinae) produce a class of defensive toxins called cardiotonic steroids (CTS) that they use to deter potential predators ^1–6^. In contrast, fireflies in the genus *Photuris* (family Lampyridae, subfamily Photurinae) cannot manufacture their own CTS. Instead, they acquire these toxins by preying on CTS-producing firefly species, using them as a defense for both themselves and their eggs (**Figure 1A**) ^7–9^. Although most Lampyridae are predatory, only *Photuris* is documented to frequently prey on other fireflies. Among other adaptations associated with this specialization ^10^, female *Photuris* mimic the courtship signals of other female Lampyridae (including congeners) to attract male prey, earning them the moniker “*femmes fatales*” ^7,11^.

**Figure 1.**
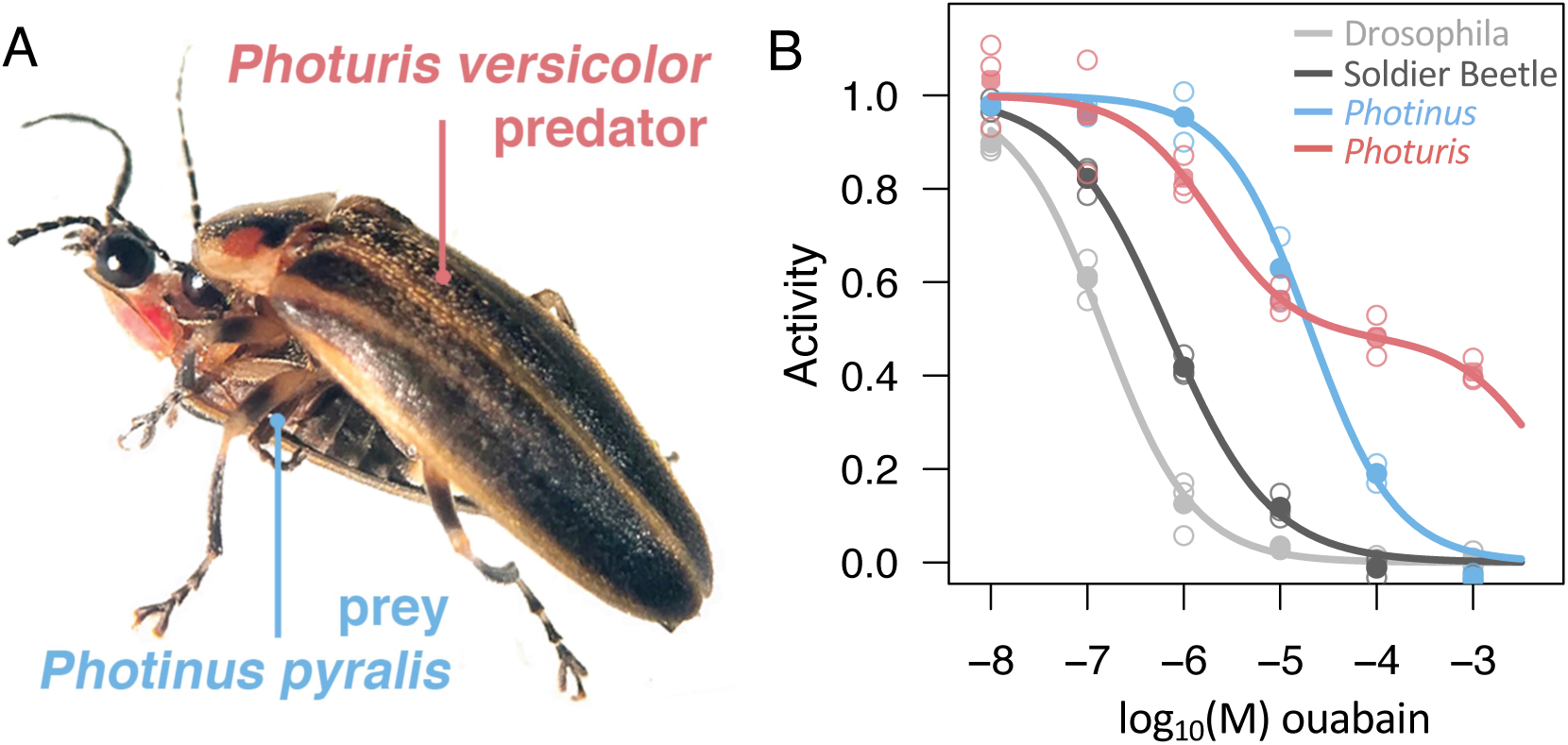
Na^+^,K^+^-ATPase (NKA) resistance to cardiotonic steroids (CTS) in predatory and prey firefly species. (A) Female *Photuris versicolor* preying on a *Photinus pyralis* male (photo by Lu Yang 2017). (B) CTS-inhibition assay on firefly brain and ventral nerve cord NKA shows that both *Photuris versicolor* (red) and *Photinus pyralis* (blue) are relatively resistant to ouabain (a representative CTS). Mean relative ATPase activity of NKA (closed circles) is plotted as a function of increasing molar (M) concentrations of the CTS ouabain. Open circles indicate biological replicates (n=3 for all except *Photinus* n=2). The NKA of both fireflies are significantly more resistant than that of the Red Soldier Beetle (*Rhagonycha fulva*) and Drosophila (*D. melanogaster* w1118) (Table S4). The fitted curves for Drosophila, Red Soldier Beetle and *Photinus pyralis* assume monophasic inhibition functions, whereas that for *Photuris versicolor* is assumed to be biphasic (see Methods).

CTS act by inhibiting Na^+^,K^+^-ATPase (NKA), an essential enzyme that helps maintain homeostasis in animals ^12,13^. CTS-resistant forms of NKA have evolved repeatedly across diverse insects and vertebrates via amino acid substitutions to the CTS-binding domain of the enzyme’s alpha-subunit (ATPα, more specifically ATPα1) ^14–22^. CTS-adaptation in insects exhibits two distinct recurrent patterns of ATPα molecular evolution. The first is that CTS resistance typically evolves via a small number of amino acid changes, and most frequently via substitutions to three sites (111, 119 and 122) in the first extracellular domain (H1-H2) of the protein ^15,22,14,23,24^. A second recurrent pattern in CTS-adapted insects is the frequent duplication and neofunctionalization of ATPα, with resistant and sensitive paralogs differentially allocated to the gut and nervous tissue ^21,22,25^. These findings generally point to a high degree of predictability in the genetic basis of CTS resistance evolution in insects ^21,22^.

Here, we investigate whether ATPα1 is also a target of CTS-adaptation in fireflies and whether these adaptations evolved via similar mechanisms in firefly predators (i.e. *Photuris*) and other firefly lineages. *Photuris* species share a number of adaptations with other fireflies including warning coloration and bioluminescent signaling (with associated structures). Collectively, these similarities likely reflect a combination of shared history and convergent evolution ^26–29^. Accordingly, it is also possible that CTS resistance in *Photuris* and other firefly species evolved either in their common ancestor or convergently via similar molecular mechanisms. Alternatively, given the different physiological challenges associated with sequestering these toxins from a food source (i.e. *Photuris*) versus producing CTS autogenously (i.e. most other firefly species) ^30–32^, it is also possible that distinct modes of CTS resistance evolved independently in *Photuris* and other firefly lineages.

## Results

### Duplication and neofunctionalization of ATPα1 in Photuris

To investigate the basis of CTS resistance in predator and prey firefly species, we first considered the level of CTS resistance of membrane-bound NKA proteins isolated from the nervous systems of wild-caught *Photinus pyralis* (a representative CTS-producing prey species), *Photuris versicolor* (a representative predator species) and compared these to the closely-related Red Soldier Beetle (*Rhagonycha fulva*) as well as Drosophila (*Drosophila melanogaster*). The NKA of both firefly species exhibit substantially higher resistance to ouabain — a water-soluble CTS — than both the Red Soldier Beetle and Drosophila enzymes (**Figure 1B**, **Table S4**). While the monophasic inhibition curves for Drosophila, the Red Soldier Beetle and *Photinus* suggest a single resistant form of the enzyme, the curve for *Photuris* is still only partially inhibited at the highest inhibitor concentrations. We hypothesized that the curve is likely to be a truncated biphasic (or multi-phasic) curve indicating that *Photuris* may have multiple isoforms of NKA that differ in their level of CTS resistance. Notably, despite having no exposure to CTS, the NKA of the Red Soldier Beetle is 4.6-fold more resistant to CTS than the Drosophila protein (**Figure 1B**, **Table S4**).

Using RNA-seq *de novo* assembly and available *de novo* genome assemblies (**Table S1**), we reconstructed ATPα1 sequences for multiple firefly species and outgroups. Consistent with our *in vitro* CTS-inhibition assays (above), we found that the Red Soldier Beetle, *Photinus* and other toxin producing firefly genera appear to have a single copy of ATPα1. In contrast, both *Photuris* species surveyed harbor multiple copies of ATPα1 that we designate as paralogs A-D (**Figure 2A**). Our analysis of RNA-seq data indicates that duplications of ATPα1 are also absent or not expressed in *Bicellonycha*, an outgroup to the *Photuris* genus (see Methods). Together with the phylogenetic tree (**Figure 2**), this evidence suggests that all three duplication events occurred after the split of *Photuris* and *Bicellonycha* (∼50 Mya ^33^), but prior to the split between *Photuris versicolor* and *Photuris frontalis.* Given the ages of these duplications relative to the *P. versicolor* and *P. frontalis* divergence (**Figure 2A**; Refs ^33,34^), it is likely that they are shared by most, if not all species of the *Photuris* genus.

**Figure 2.**
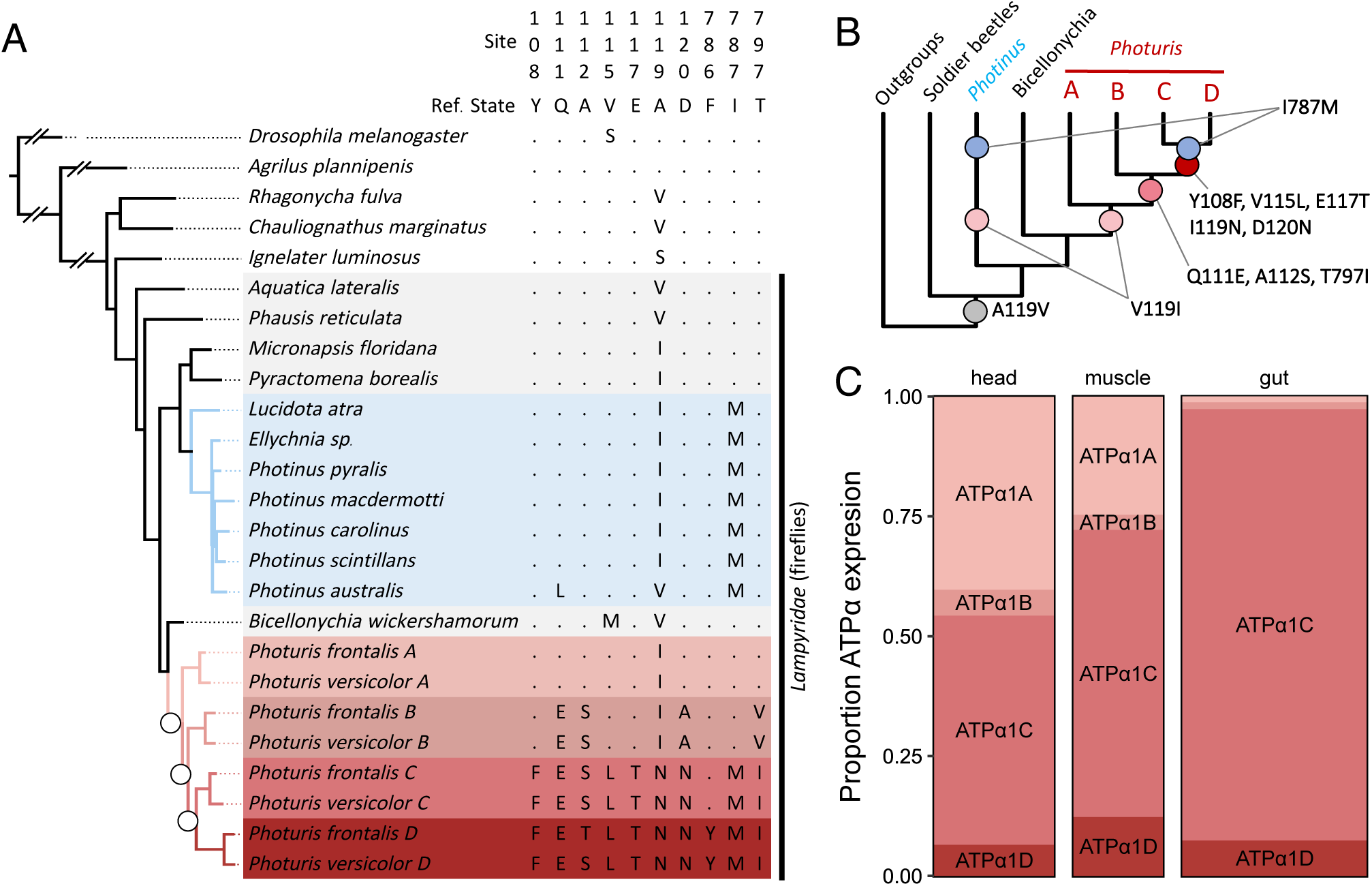
Molecular evolution of ATPα1 in fireflies. (A) A maximum likelihood genealogical tree based on ATPα1 protein-coding sequences (Methods and Supplementary Figure S1 for more detail). All species are beetles (Coleoptera) except for *Drosophila melanogaster*. CTS-producing firefly species are shaded in blue and four *Photuris* ATPα1 paralog lineages A-D are indicated in increasingly darker shades of red. White circles indicate three sequential rounds of duplication in the Photuris lineage. Substitution patterns are shown only for sites with known roles in CTS-resistance, except for site 787, which is newly discovered in this study. The reference sequence corresponds to the reconstructed ancestral sequence for beetles. Dots indicate identity with the ancestral state and letters represent derived amino acid substitutions. The phylogenetic position of *Bicellonychia wickershamorum* has 86% bootstrap support and is consistent with phylogenetic reconstructions based on multiple loci ^33,50^. (B) Schematic graph showing key amino acid substitutions associated with CTS-resistance inferred to have occurred in species and paralog lineages. Soldier beetles are represented by *Rhagnonycha* and *Chauliognathus*. See Figure S1 for more details. (C) Tissue-specific expression of ATPα1 paralogs in *Photuris versicolor* females (see also Table S2, Supplementary Figure S2). Column width corresponds to relative proportion of total normalized ATPα1 expression summed across paralogs in *Photuris*, and shaded segments to relative levels of expression of the four ATPα1 paralogs in each tissue.

*Photuris* ATPα1 paralogs are differentially expressed among tissues. Based on the pattern of amino acid substitution at sites previously implicated in CTS-resistance, ATPα1A is predicted to be the most CTS-sensitive paralog and is more highly-expressed in head relative to gut tissue (adjusted p=3.0e-6). Conversely, ATPα1C and ATPα1D are predicted to be the most CTS-resistant paralogs because they have accumulated multiple amino acid substitutions implicated in CTS resistance (**Figure 2A-B**). In contrast to ATPα1A, ATPα1C is substantially more highly expressed in the gut relative to the head (adjusted p = 8.8e-4, **Table S2**, **Figure 2C**). This implies that ATPα1 paralogs have neofunctionalized, as observed in multiple CTS-adapted insects carrying ATPα1 duplications ^21,22,25^. ATPα1A and ATPα1C together comprise 88% of ATPα1 transcripts in the female *Photuris* head and are represented in roughly equal proportions (**Figure 2C**), lending support to the “biphasic” interpretation of the enzyme-inhibition curve for *Photuris* nervous tissue (**Figure 1B**). Given the stepwise accumulation of amino acid substitutions among *Photuris* paralogs (**Figure 2B**), we used CRISPR-Cas9 genome-editing (see Methods), and a similar site-directed cassette exchange system^23^, to generate a series of *Drosophila melanogaster* strains that carry substitutions occurring at key stages of ATPα1 neofunctionalization in *Photuris* (**Figure 3A**).

**Figure 3.**
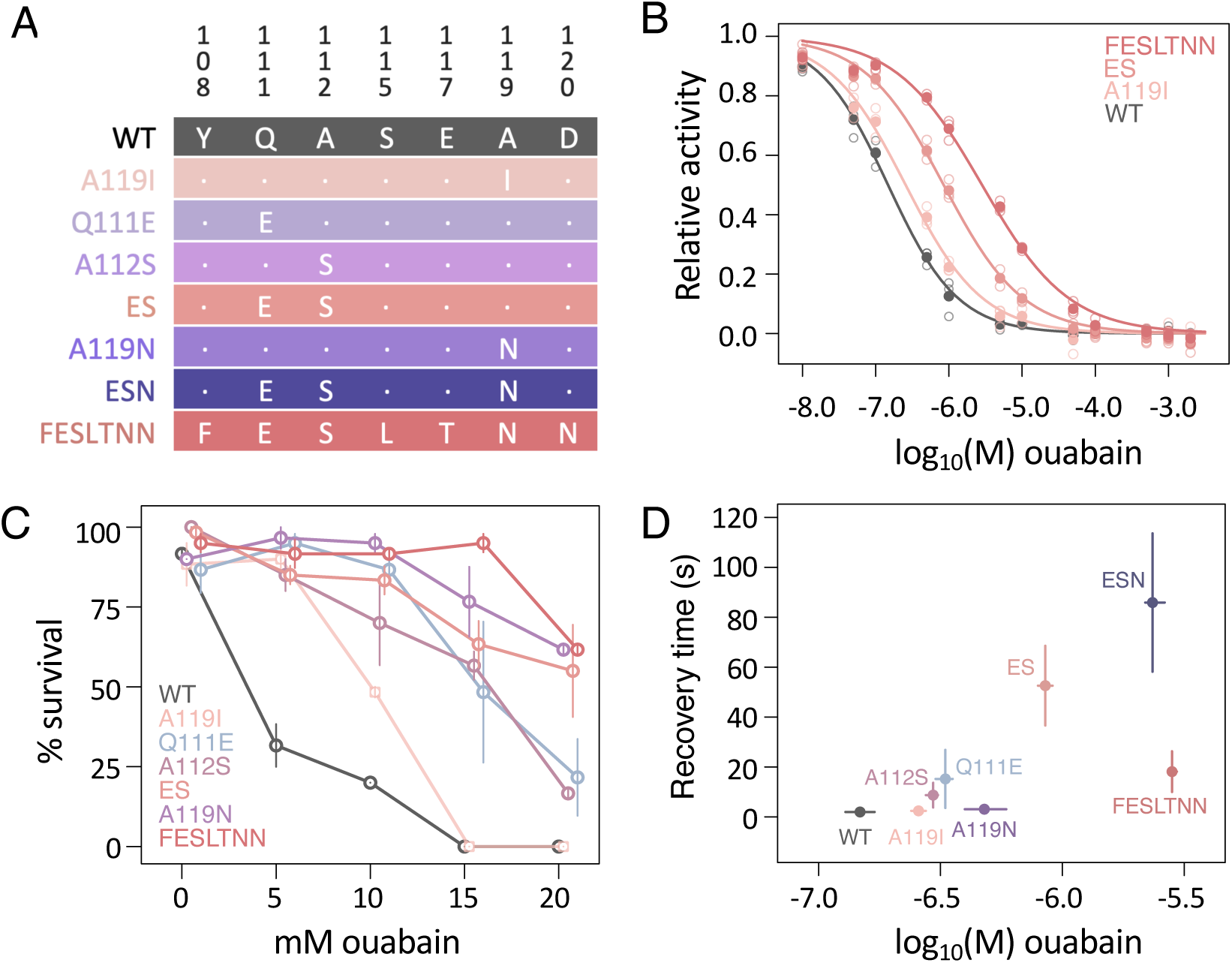
Functional effects of *Photuris* paralog-specific substitutions. (A) Engineered fly lines generated in this study. Only sites implicated in CTS-resistance are shown. States for the CTS-sensitive *D. melanogaster* strain (w^1118^) used to generate engineered lines are denoted “WT” for “wild-type”. All strains were homozygous for the assayed substitutions. Detailed results for additional engineered lines can be found in Supplementary Figures S3 and S4. (B) CTS-inhibition assays for NKA isolated from heads of WT and engineered fly lines. Mean relative activities (filled circles) are plotted as a function of increasing concentrations of ouabain, a representative CTS. Each mean is the average of three biological replicates. Solid lines represent the least squares fit model (as per Figure 1; Methods). (C) CTS-tolerance of wild-type and engineered adult flies. Survival of adult flies upon seven days exposure to increasing concentrations of ouabain. Points represent the average of three biological replicates (pools of n=20 individuals), and whiskers correspond to standard errors. “ESN” could not be assayed here due to the poor condition of the line. (D) Levels of neural dysfunction and enzyme resistance to ouabain for wild-type (WT) and engineered fly lines. The neural dysfunction assay (y-axis) measures the recovery time (seconds) following seizures induced by mechanical overstimulation (aka the “bang sensitivity” assay) of adult male flies. 28-43 individuals (open circles) are assayed for each line. The level of enzyme resistance to ouabain (x-axis) is measured as IC50 in enzyme-inhibition assays (as per Figure 1; Methods). Plotted are the means and 95% bootstrap confidence bounds as points and whiskers, respectively.

### Stepwise accumulation of CTS resistance substitutions in firefly lineages

Phylogenetic inference reveals that the first steps in the evolution of ATPα1 CTS-resistance were shared between *Photuris* and other firefly lineages. Specifically, the substitution of alanine to valine at position 119 (A119V) likely preceded the origin of fireflies (**Figure 2B**; **Supplementary Figure S1**). The higher CTS resistance of Red Soldier Beetle NKA relative to Drosophila (**Figure 1B**) is consistent with a contribution of this substitution to NKA CTS-resistance. A subsequent substitution to the same site, V119I, is observed in both *Photuris* and other firefly species. *Bicellonycha* is a member of a sister genus to *Photuris* ^33^ and the species we surveyed (*B. wickershamorum*) lacks V119I. In fact, our phylogenetic analysis suggests that *Bicellonycha* retains the ancestral state and that V119I most likely evolved convergently in both *Photuris* and other firefly species soon after they became distinct lineages (**Figure 2B**; **Supplementary Figure S1**).

To estimate the functional effects of the eventual transition from alanine to isoleucine at position 119 in *Photuris* and other firefly lineages (i.e. A119I = 119A -> 119V -> 119I), we modified the *D. melanogaster* ATPα1 protein using genome-editing tools. We find that A119I modestly but significantly increases CTS resistance of NKA relative to the sensitive wild-type *D. melanogaster* enzyme (1.8-fold, 95% CI: 1.6-1.9; **Figure 3B**). A119I also substantially improves *D. melanogaster* adult survival upon exposure to CTS (**Figure 3C**). Since NKA is critical for proper neural function, we subjected adult engineered flies to mechanical over-stimulation and measured recovery time from induced seizures (the so-called “bang sensitivity” assay, ref. ^35^, see Methods). We found that A119I mutant flies exhibit no obvious neurological dysfunction compared to wild-type flies (**Figure 3D**). Taken together, we conclude that the evolution of alanine to isoleucine at position 119 comprises a potential exaptation that may have facilitated the emergence of both the ability of fireflies to manufacture CTS and the predatory specialization of *Photuris*.

Despite these shared early steps in the evolution ATPα1 CTS-resistance, it is apparent that lineages leading to the predatory genus *Photuris* subsequently took a radically different approach compared to other fireflies: repeated duplication and neofunctionlization of ATPα1. Following an initial duplication of ATPα1 in the *Photuris* lineage, two paralogous lineages began to diverge in function, with one lineage ancestral to more resistant paralogs B-D. The most conspicuous candidate CTS resistance substitutions on the B-D lineage are Q111E and T797I (**Figure 2B**). Site 111 is a known hotspot for convergent CTS resistance substitutions in animals ^14,17,22^. At site 797, the ancestral threonine residue is predicted to form a stabilizing hydrogen bond between ATPα1 and CTS ^36^ that the derived T797I substitution (along the paralog B-D lineage) is predicted to disrupt. Previous work established that the biochemically similar substitution T797V results in an 80-fold increase in CTS resistance of mammalian NKA^37^. A substitution to a third site in the paralog B-D lineage, A112S, was also of interest as this site was previously identified as a target of positive selection in CTS-resistant toads and their predators ^18^. Further, A112S repeatedly co-occurs with Q111E/R/T across phylogenetically diverse taxa including both insects and vertebrates ^17,23^.

Given these substitution patterns, we engineered a series of *D. melanogaster* lines to dissect the basis for CTS resistance along the lineage leading to paralogs B-D. We began by focusing on substitutions at sites 111 and 112. NKA isolated from fly lines engineered with Q111E+A112S (hereafter “ES”) exhibits a 6-fold increase in CTS resistance relative to the wild-type enzyme of Drosophila (**Figure 3B**; **Table S4**). Together, these substitutions also confer engineered adult flies with substantial levels of resistance to CTS exposure (**Figure 3C**, **Table S4**). Q111E and A112S each have significant effects individually (both ∼2-fold) and their combined effects appear to be close to additive (**Supplementary Figure S3**; **Table S4**). We also find that the substitutions Q111E and A112S cause slight neural dysfunction when introduced individually, and this dysfunction is exaggerated when they are combined (in “ES” flies, **Figure 3D**, **Supplementary Figure S3**). These results suggest that the substantial CTS resistance conferred by this combination is associated with negative pleiotropic effects on protein function. In *Photuris* these may be compensated either by other substitutions in ATPα1 paralogs B-D or by other mechanisms.

While we did not generate lines representing the full complement of evolutionary paths along the paralogs B-D lineage, we find that substitution of T797I is homozygous lethal on the *D. melanogaster* ATPα1 background. This is not unexpected given previous work showing that the similar substitution, T797V, decreases NKA activity to 3.4% of wild-type levels ^37^. Interestingly, the isoleucine at site 797 of *Photuris* ATPα1B was replaced with valine after a second round of duplication (i.e. I797V, **Supplementary Figure S1**). Additional engineering reveals that the combination Q111E+A112S+A119I+D120A (hereafter “ESIA”), occurring along the ATPα1B lineage, is also homozygous lethal. It may be that detrimental effects of ESIA and T797I (and subsequently I797V) observed on the *D. melanogaster* protein are ameliorated when combined (i.e. ESIAI/V) or by other substitutions in the ATPα1 sequence background of fireflies. However, even if active and resistant, ATPα1B has the lowest expression among the four paralogs (**Figure 2C**), implying that it may contribute little to overall CTS resistance in *Photuris*.

A more dramatic series of substitutions implicated in CTS resistance occurs along the lineage leading to *Photuris* ATPα1 paralogs C and D (**Figure 2B**). We engineered most of these CTS-relevant substitutions (Y108F+Q111E+A112S+V115L+E117T+A119N+D120N, hereafter “FESLTNN”) into the *D. melanogaster* ATPα1 protein. The FESLTNN combination results in an additional 3-fold increase in NKA CTS-resistance over ES alone **(Figure 3B; Table S4)**. FESLTNN adults also exhibit exceptionally high rates of survival upon CTS exposure (**Figure 3C)**. Further dissection of individual substitution effects reveals that A119N and the combination (Q111E+A112S+A119N, hereafter “ESN”) confer substantial CTS resistance to NKA (almost 3-fold and 16-fold, respectively). The level of NKA CTS resistance for ESN (16-fold) is only slightly lower than for FESLTNN (18-fold, **Table S4**), implying that the marginal effect of A119N is nearly sufficient to explain the difference between ES and FESLTNN. A119N alone also appears to be sufficient to confer levels of adult resistance to CTS exposure that are comparable to FESLTNN (**Figure 3; Supplementary Figure S4C**).

Evaluation of the trade-offs associated with resistance conferred by various evolutionary intermediates offers further insight into the likely evolutionary paths used to evolve CTS resistance along the C/D paralog lineage (**Figure 3D**; **Supplementary Figure S4**). Notably, “ES” confers substantial CTS-resistance to the *D. melanogaster* protein, but at the cost of neurological defects (**Figure 3D**). While adding A119N to the ES background (i.e., ESN) results in even higher CTS resistance, this comes at the cost of even greater neurological dysfunction. Interestingly, we find that ESN in combination with other substitutions added to the C/D paralog lineage (i.e. FESLTNN) results in a similar level of CTS resistance, but with substantially reduced neurological dysfunction. Taken together, our results reveal that serial rounds of gene duplication and neofunctionalization led to stepwise increases in the level of CTS resistance in *Photuris* over time with substitutions A119I, Q111E+A112S and I119N being key evolutionary steps. In addition, substitutions at other sites in *Photuris* NKAs, while not contributing much to resistance itself, nonetheless appear to play a critical role in ameliorating pleiotropic costs associated with key resistance substitutions. Similar patterns of background dependence (aka “intra-molecular epistasis “) have been observed in the evolution of CTS-resistant forms of NKA in insects and vertebrates^23,24,16,17,38^, the evolution of nicotinic acetylcholine receptor resistance to epibatidine in dendrobatid frogs ^39^ and γ-aminobytyric acid (GABA) receptor resistance to the insecticide fipronil in plant hoppers ^40^.

### The evolution of CTS resistance in prey firefly species

The *Photinus* NKA protein is also highly resistant to CTS inhibition *in vitro* (**Figure 1B**). However, in contrast to the dramatic patterns of neofunctionalization in *Photuris*, the sole copy of ATPα1 in *Photinus* curiously lacks substitutions at known CTS-insensitivity sites other than V119I (**Figure 2**; **Supplementary Figure S1**), which has a significant but relatively small effect (**Figure 2B, Table S4**). Notably absent in prey firefly species are substitutions at sites 111 and 122 which are most often associated with large effects on CTS resistance in species with a single copy of ATPα1 ^21,22^.

In order to search for previously undocumented CTS-resistance sites throughout the protein, we queried all substitutions, naive to previously known functional importance, in a large alignment of ATPα1 sequences from CTS-adapted and non-adapted insects (see Methods). Of potential candidate substitutions in the *Photinus* lineage, only one (I787M) exhibits a strong signature of parallel evolution in multiple CTS-adapted taxa (**Figure 4A; Supplementary Figure S5**). Notably, I787M appears to be present in all CTS-producing firefly species surveyed here (*Lucidota*, *Ellychnia*, *Photinus*), and is also present in paralogs C and D of *Photuris* (**Supplementary Figure S1**). Using CRISPR-Cas9 engineering of *D. melanogaster*, we show that I787M has a significant effect on CTS resistance of the enzyme *in vitro* (a 3-fold increase; **Figure 4B, Table S4**). Despite its relatively modest effect on CTS-resistance of the enzyme *in vitro*, I787M has a substantial effect on the tolerance of adult *D. melanogaster* to CTS exposure (**Figure 4C**) and is not associated with substantial neurological dysfunction (**Figure 4D**). Thus, I787M is a previously unreported determinant of CTS-resistance that shows a high degree of parallelism among CTS-adapted insects.

**Figure 4.**
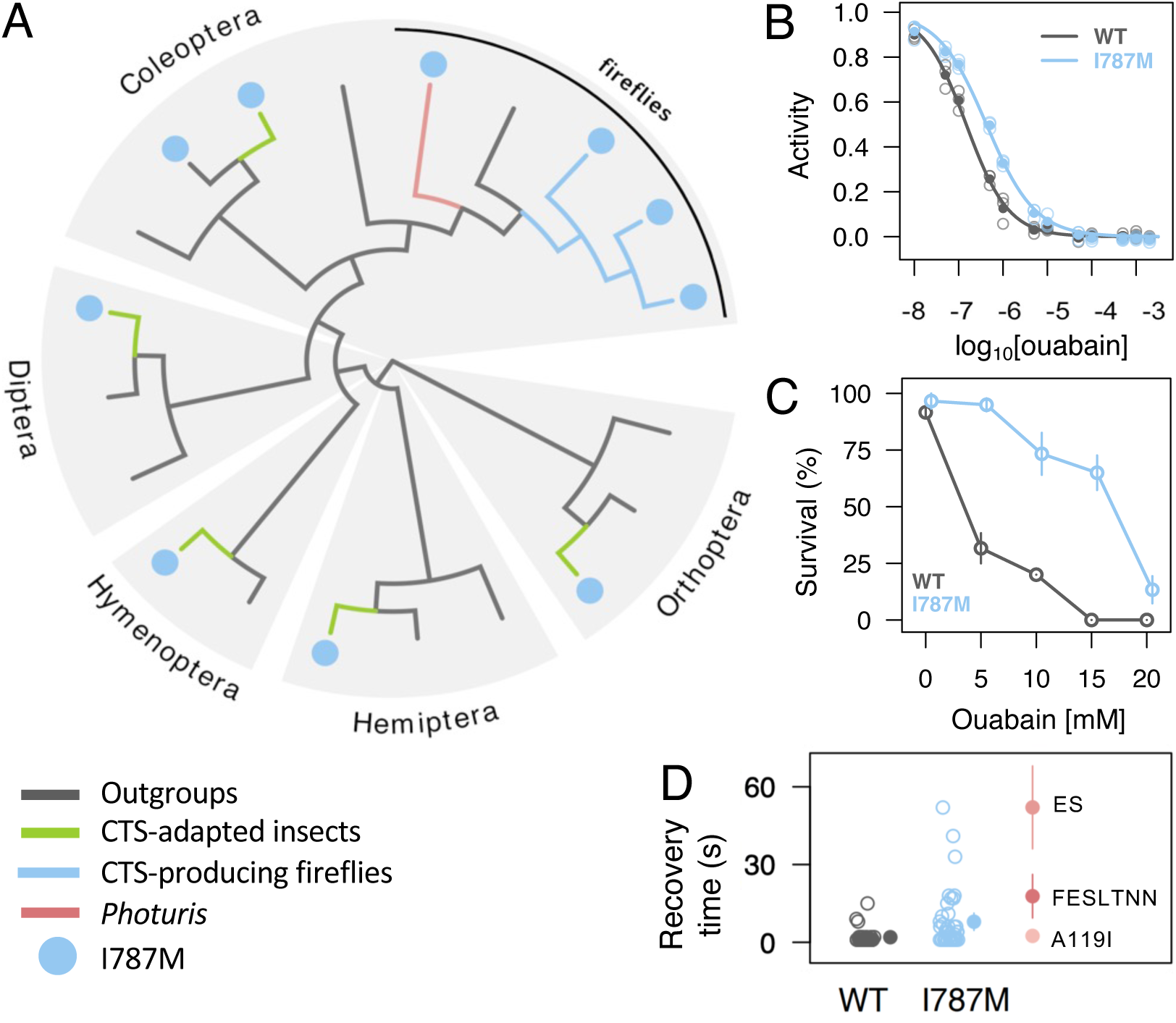
Evolutionary and functional analysis reveals a new CTS-resistance substitution, I787M. (A) Recurrent appearance of I787M in CTS-associated herbivores (green lineages), CTS-producing prey fireflies (blue lineages) and predatory *Photuris* fireflies (red lineage). See Supplementary Figure S5 for more details. (B) Estimates of the IC50 of NKA of wild-type and engineered fly lines in CTS-inhibition assays. The log_10_(IC_50_) of I787M flies is 2.9-fold higher than that of WT (*w*^1118^) control flies (Table S4). Each estimate is based on three biological replicates (each with three technical replicates). (C) CTS-tolerance of adult wild-type and engineered flies. Engineered flies carrying I787M exhibit higher survival rates upon seven-day CTS exposure. (D) Neural dysfunction as measured by the “bang sensitivity” assay. The assay measures the time to recovery (seconds) from seizures induced by mechanical over-stimulation (as in Figure 3C). Open circles correspond to individual flies. Filled circles to the right are means for ES, FESLTNN, and A119I with 95% CI whiskers (individual data points not shown). Note that for ES, FESLTNN, and A119I, individual data points are not shown. I787M individuals are somewhat more susceptible to mechanical over-stimulation than WT (Wilcoxon test p= 3.4e-5), but the impairment is less severe than observed for some other engineered strains. See Figure 3D for more details on the A119I, ES, FESLTNN lines.

## Discussion

CTS resistance and the use of CTS for defense have convergently evolved in a large number of species. Our study reveals that the first steps toward CTS resistance evolution in fireflies (A119V, V119I) were likely taken before CTS synthesis evolved in *Photinus* and before predatory specialization on fireflies emerged in *Photuris*. One possible explanation for these shared steps is that *de novo* production of CTS is ancestral to fireflies and that the ability to do this was subsequently lost in *Photuris* as they opted for predation as an alternative source of these toxins. However, there is little evidence for this based on the phylogenetic distribution of CTS production in fireflies (**Figure 4A** and see Ref. ^1^). An alternative hypothesis is that these steps may reflect exaptations in fireflies unrelated to CTS metabolism. This is supported by the widespread distribution of substitutions at site 119 among insect taxa that are not adapted to CTS^23^, including the Red Soldier Beetle surveyed here, which is closely related to fireflies. Interestingly, A119N – a key substitution underlying resistance of neofunctionalized *Photuris* paralogs – is present in all Hymenopteran species surveyed to date^23^. As most Hymenopterans are not associated with CTS, A119N may function as an exaptation that facilitated herbivorous wasp specialization on CTS containing hostplants ^41^, parasitoid wasp species predation on CTS-adapted herbivorous insects ^42^ or as generalist nectar feeders on the flowers of CTS-producing plants ^43^.

Despite initial shared steps toward CTS resistance taken early in firefly diversification, our study also highlights the distinct dynamics of CTS resistance evolution associated with their different ecological roles. Although both *Photuris* and other fireflies are protected by the same class of CTS toxins, they nonetheless face different biochemical and physiological challenges posed by *de novo* production (most firefly genera) versus sequestration from a food source (*Photuris*) ^30–32^. Notably, the dramatic repeated duplication and neofunctionalization of ATPα1 in *Photuris* resembles the strategy employed by several CTS-adapted herbivores that sequester CTS from a food-source ^19,21,22,25^. We speculate that the duplication and neofunctionalization of ATPα1 in *Photuris* may have been driven by the evolution of firefly predation in the *Photuris* lineage, which imposed similar biochemical and physiological challenges to those experienced by herbivores that sequester CTS from hostplants.

We have shown that intermediate stages in the evolution of resistant forms of NKA in *Photuris* are associated with substantial enzymatic and physiological dysfunction when engineered into *D. melanogaster*. This dysfunction would likely represent a substantial negatively pleiotropic barrier to the evolution of CTS resistance if the ATPα1 gene existed as a single copy in *Photuris*. The role of gene duplication as a solution to overcoming pleiotropic constraints associated with intermediate states has previously been proposed for genes involved in Galactose metabolism in yeast ^44^, ribonuclease genes in primates ^45^ and the evolution of lens transparency in the eyes of vertebrates ^46^, among other examples. We propose that these negative pleiotropic effects were largely avoided by duplication and neofunctionalization in the *Photuris* lineage, allowing this lineage to specialize on consuming and sequestering toxins from CTS-containing prey. Thus, our work implicates duplication and neofunctionalization as a potential factor in the diversification of species roles within an ecological community.

There is a strong association in insects between duplication and neofunctionalization of ATPα1 and sequestering CTS from a food source. Specifically, ATPα1 in *Photuris* currently represents the 6^th^ documented case with no counter-examples of duplications observed in non-sequestering species^21–23^. Despite this, there are several clear examples of species that sequester high levels of CTS from a food source that lack neofunctionalized ATPα1 duplications (for e.g. the Monarch butterfly). A related question is why CTS-producing firefly species lack duplication and neofunctionalization of ATPα1 despite sequestering high levels of CTS. The reasons for this difference may be manifold. First, while CTS-producing firefly species indeed store CTS, they produce these compounds internally and do not absorb them from a food source. Additionally, even insect species that are not adapted to CTS exhibit physiological features that at least partially protect them from the adverse effects of CTS exposure, including restricted expression of NKA to neurons (in Lepidoptera^47^) or expression of protective proteins that buffer the adverse effects of dietary CTS (for e.g. in Lepidoptera^47^; in Drosophila^48^). It is likely that the necessity of duplication and neofunctionalization of ATPα1 depends on factors such as the amount of CTS typically consumed, the polarity of these CTSs, the permeability of tissues and cells to CTS, among other factors. Further research is needed to determine which, if any, of these factors account for patterns of duplication and neofunctionalization of ATPα1 in fireflies and other CTS-adapted insect species.

There remain some interesting and puzzling differences between insect herbivores and fireflies with respect to the molecular basis of CTS-resistance via substitutions to ATPα1. A recent study showed that three amino acid substitutions are sufficient to account for the entire difference in NKA CTS-resistance observed between the wild-type proteins of *D. melanogaster* and the monarch butterfly, *Danaus plexippus* ^24^. In contrast, our attempts to engineer known CTS-resistance substitutions of fireflies into *D. melanogaster* NKA fall short of the level of CTS-resistance level of wild-type firefly proteins (**Figure 5**). This suggests wild-type firefly NKAs achieve high levels of CTS-resistance via substitutions at other sites in ATPα1. Previous attempts to map determinants of CTS resistance using saturation mutagenesis are likely to have missed sites with relatively small effects on CTS resistance ^49^. This may explain why the I787M, discovered here using phylogenetic methods, was missed in previous screens for CTS-resistance mutations. The gap between engineered-*Drosophila* and wild-type firefly NKA CTS-resistance, and the lack of obvious candidate substitutions in *Photinus*, suggests that there are likely to be other, as yet undiscovered, determinants of CTS resistance in firefly NKAs.

**Figure 5.**
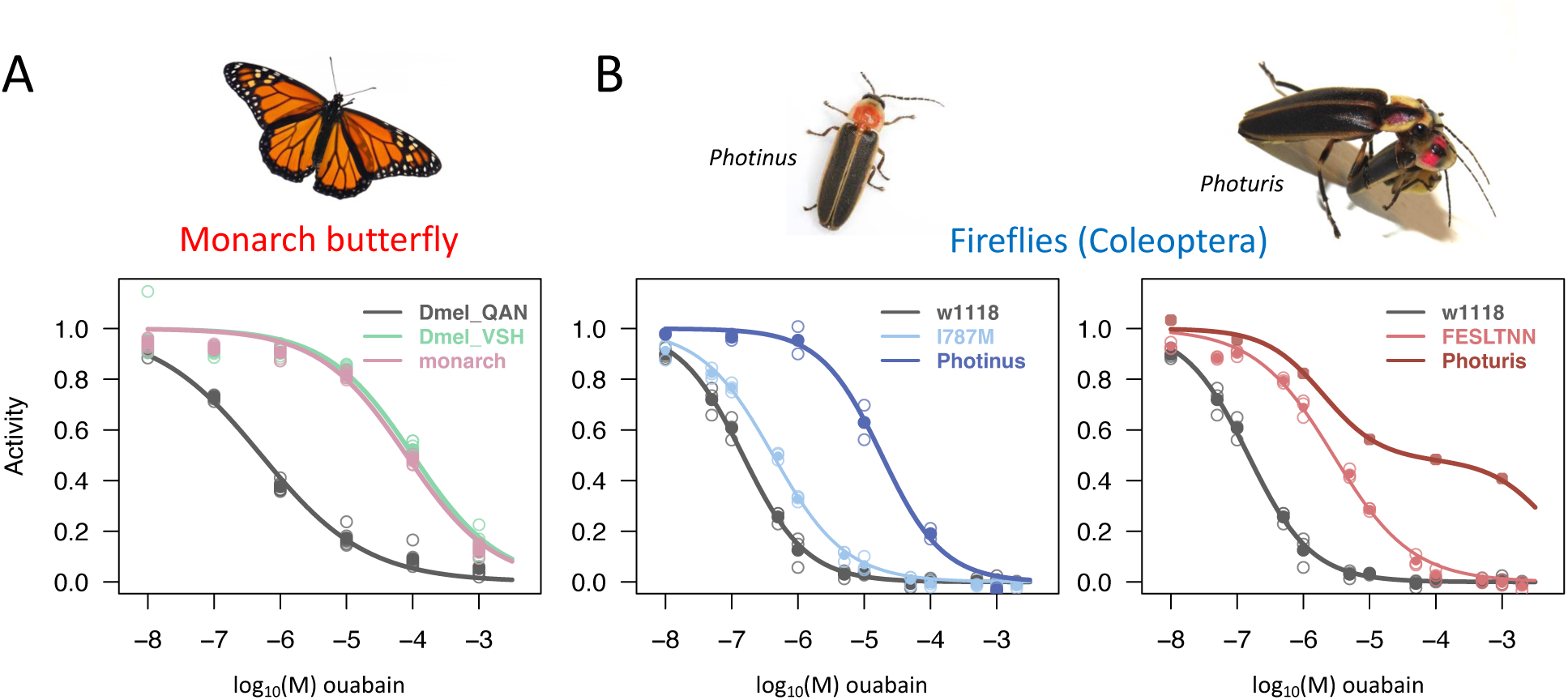
CTS-inhibition curves suggest as yet unmapped determinants of CTS resistance in firefly Na^+^,K^+^-ATPases (NKAs). (A) Engineering three known CTS resistance-associated amino acid substitutions into *D. melanogaster* NKA accounts for ∼100% of the CTS resistance of the Monarch butterfly protein (data from Karageorgi et al. 2019). Plots are as in Figure 1 of main text. Dmel_QAN = wild type *D. melanogaster* protein; Dmel_VSH = *D. melanogaster* protein + substitutions Q111V, A119S and N122H; monarch = Monarch butterfly (*Danaus plexippus*) protein. In contrast, engineering the known major CTS resistance substitutions observed in fireflies into *D. melanogaster* NKA results in only a fraction of the observed CTS resistance of the *Photinus* (left panel) and *Photuris* (right panel) proteins. The latter pattern points to as yet unmapped determinants of CTS resistance of firefly NKAs. w1118 = wild-type *D. melanogaster*. The I787M and FESLTNN constructs are described in Figures 4 and 3, respectively. “Photinus” and “Photuris” are as in Figure 1.

## Acknowledgements

We thank the Fireflyers International Network for providing samples of *P. frontalis* (notably H. Morgan and P. Shaw) and their valuable natural history input. We thank P. Reilly for advice on CRISPR-cas9 design, A. Taverner for help with fly crosses, C. Han for advice on variant calling analysis, and M. Kardestuncer for help with collections. We thank S. Stiehler and M. Jang for technical assistance. Finally, we thank AM for inspiration. This work was funded by the National Institutes of Health R01 GM115523 to PA, AG059385 to MJP, and DFG PE2059/3-1 and the LOEWE program of the State of Hesse (Insect Biotechnology & Bioresources), Germany to GP.

## Author contributions

PA conceptualized the study; LY, FB, MA, RV, MW and PA collected and prepared firefly samples; LY, FB, YZ, MA and JP generated sequence data; LY, FB and PA performed data analysis; MJP, BPR and ADT generated phiC31/cre-lox engineered flies. LY generated a subset of CRISPR/cas9 engineered flies; LY, FB, AB, GP and PA designed and performed functional assays; Funding acquisition and supervision by PA, MJP and GP. PA wrote the paper with input from the other authors.

## STAR Methods

### Resource Availability

#### Lead Contact

Further information and requests on methods can be directed to Dr. Peter Andolfatto

#### Materials Availability

Plasmids used in this study are available upon request. This study did not generate new unique reagents.

#### Data and Code Availability

Data generated during this study are available through links provided in the Key Resources Table. Nucleotide sequences (CDS) of ATPα1 have been submitted to GenBank (Accession Numbers: MT897473-MT897482). RNA-seq data for *Photinus pyralis* and *Photuris versicolor* have been deposited in the Sequence Read Archive (BioProject PRJNA891248 and PRJNA891306/ PRJNA922528, respectively). This paper does not report original code. Any additional information required to reanalyze the data reported in this work paper is available from the lead contact upon request.

### Experimental Model and Study Participants Details

Wild specimens of adult *Photinus pyralis* and *Photuris versicolor* were used. The origin of all *Drosophila* fly strains used can be found in the Key Resources Table. All flies were cultured on standard cornmeal-agar medium in uncrowded conditions unless stated in the methods.

### Quantification and Statistical Analysis

To determine the difference of expression level of NKA among tissues and paralogs, we used the beta-binomial (ibb) test^51^ followed by a standard Bonferroni correction. Statistical tests are summarized in **Table S2**. To evaluate differences in adult tolerance to CTS exposure, the Cochran-Mantel-Haenzel test implemented in R was used to assess significant differences between ouabain treatments (5/10/15/20 mM) versus no ouabain. We used 3 replicates (20 individuals each) per line per condition. Mean and standard errors are represented Figure 3 and 4. To determine the relative activities of NKA for each ouabain concentration, 95% confidence intervals of IC50 were estimated by parametric bootstrapping as described in ref. ^23^. Estimated parameters can be found in **Table S4**. Number of biological replicates used is indicated in the legends of **Figure 1**, **Figure 3** and **Figure 4**. For bang-sensitivity assays, 30-50 individuals were tested for each genotype. Mean and standard errors are represented in **Figures 3 and 4**.

### Method Details

#### Firefly collection and data sources

*Photuris versicolor* and *Photinus pyralis* were collected in Princeton, NJ, USA during the summer months of 2017, 2018 and 2022. Specimens of *Rhagonycha fulva* were collected in Stuttgart, Germany in July 2023. Samples were immediately stored at −80°C with or without RNAlater (InvitrogenTM) depending on the planned downstream experiments. For other species, we used publicly available data (see **Table S1**).

#### RNA-seq data generation and ATPα1 reconstruction

Total RNA from muscle (thoracic or leg), heads and gut of male *Photinus*, female and male *Photuris* fireflies was extracted with TRIzol (Ambion, Life Technologies) following the manufacturer’s protocol. For male *Photinus* and female *Photuris* samples, RNA-seq libraries were prepared with TruSeq RNA Library Prep Kit v2 (Illumina) and sequenced on a HiSeq4000 (Genewiz, South Plainfield, NJ, USA). Reads were trimmed for quality and length using TQSfastq.py (Key Resources Table) with default parameters. For male *Photuris* samples, RNA-seq libraries were prepared with TruSeq Stranded mRNA Library Prep (Illumina) and sequenced on HiSeq X (PSOMAGEN, Inc.). Reads were trimmed for adapters but not for quality using trim_galore (v0.6.7) with cutadapt (v1.18). The three RNA-seq datasets were used to generate *de novo* transcriptome assemblies using Trinity v2.2.0^64^ with default parameters. Beetle ATPα1 sequences^22^ were used as initial queries to BLAST (BLAST 2.13.0+ with default parameters) against firefly transcriptome assemblies. Reconstructed ATPα1 sequences for *Photinus pyralis* and *Photuris versicolor* were then used as templates to identify ATPα1 sequences in *de novo* transcriptome and genome assemblies from other firefly species.

Since *de novo* assembly of *P. versicolor* revealed four distinct paralogs of ATPα1, we set out to determine the ages of duplication events by surveying closely related species *Photuris frontalis* and *Bicellonycha wickershamorum* (**Table S1**). We searched for ATPα1 homologs in the *de novo* transcriptomes of *P. frontalis* and *B. wickershamorum* using BLAST (tblastn, BLAST 2.13.0+, evalue=1e-50). Orthologs of all four *P. versicolor* paralogs we unambiguously identified in *P. frontalis.* To reconstruct the *P. frontalis* paralogs, RNAseq reads were mapped to *P. versicolor* paralogs using bwa mem (v 0.7.17) as described above. Duplicates were identified and reads were assigned a read-group using Picard (v 2.27.5). BAM files were sorted and indexed using Samtools (v 1.6). Reads were then realigned using GATK3 (v 3.8.1) and variants were called using the YourePrettyGood pseudoreference pipeline (Key Resources Table; samtoolsVariantCall.sh and vcfToPseudoref.sh with thresholds MQ <=20, QUAL<=26). bcftools (v1.9) was then used to create an inferred sequence for each paralog.

In contrast to *P. frontalis*, we detected only one copy of ATPα1 in the *B. wickershamorum de novo* transcriptome. The top BLAST hit shares 97.7% amino acid identity with the *P. versicolor* paralog A (score= 1882, evalue=0.0). The second-best hit (score=326, evalue=3e-94) shares only ∼25% amino acid identity with *P. versicolor* ATPα1 paralogs. The latter protein is most likely to be sarco/endoplasmic reticulum-type Ca^2+^-ATPase based on comparisons to the *D. melanogaster* genome annotation (with which it has 80% amino acid sequence identity). In a second attempt to detect possible duplications in *B. wickershamorum*, we mapped RNAseq reads to its ATPα1 and looked for potential amino acid variants supported by three or more reads. Only one amino acid variant was found at position 830 (D/E), but this site has not been implicated in CTS resistance. These lines of evidence suggest that *B. wickershamorum* lacks neofunctionalized duplications ATPα1, or that they are not expressed.

#### Confirmation of ATPα1 duplicates in *P. versiciolor*

The four reconstructed ATPα1 paralogs (A-D) of *Photuris versicolor* were confirmed by PCR-cloning and sequencing (see Key Resources Table for primers used). ATPα1 paralogs were amplified from cDNA using Phusion High-Fidelity DNA Polymerase (Thermo Fisher Scientific) and separated by agarose gel electrophoresis. The appropriate bands were gel-extracted and cleaned with QIAquick PCR Purification Kit (Qiagen). Purified PCR products were 3′A-tailed using Taq polymerase (NEB) and cloned into TOPO TA-cloning vector (Invitrogen). Plasmids with inserts were identified and isolated using colony-PCR. Tn5-tagmentation libraries^52^ were prepared for each plasmid and indexed using customized Illumina-style i7 and i5 PCR primers added with 10 cycles of PCR. The libraries were pooled and sequenced with 150 nt, paired-end reads on an Illumina MiSeq Nano flowcell. 10,000 pair-end reads per plasmid were randomly sampled, trimmed for quality and *de novo* assembled using Velvet^53^ and Oases^54^. The sequences for each paralog were aligned and visualized in SeaView^55^.

#### Phylogeny estimation and reconstruction of ancestral states

Firefly ATPα1 protein-coding sequences together with ATPα1 sequences of *D. melanogaster*, *Agrilus planipennis* and the soldier beetles *Chauliognathus marginatus* and *Rhagonycha fulva* (**Table S1**) were aligned using MUSCLE (v 3.8.425). The sequences were trimmed to produce an alignment with no gaps or missing data. A phylogenetic tree for ATPα1 sequences was estimated using maximum likelihood with PhyML (v 3.3.20180621) with the GTR model and default parameters. Ancestral sequences and substitutions along specific-lineages were estimated based on this alignment and the tree using PAML’s baseml function (v 4.9) with the following parameters: model=7, kappa=1.6, RateAncestor=2^56^.

To look for substitutions that occur recurrently in CTS-associated species, we combined the multispecies alignment of Taverner et al.^23^, which includes 174 predicted ATPα1 sequences from 161 insect species, with our alignment for firefly species (above). Considering all amino acid substitutions, naive to functional importance, we looked for those that 1) are present in at least 3 out of 7 *Photinus* species, 2) are not shared with direct outgroups, e.g., *Pyractomena*, 3) are common in other CTS-associated species relative to other species. One site, and one substitution in particular (I787M) passes these filter criteria. I787M has independently evolved in five insect orders, and 6 out of 7 times it occurred in insects that either sequester or produce CTS (**Supplementary Figure S5**).

#### Differential expression analysis

*Photuris versicolor* females were fed with *Photinus pyralis* one day before dissection. RNA-seq data was generated as described in earlier sections. We created a modified *Photuris de novo* transcriptome reference by first identifying and removing any BLAST hits matching our reconstructed ATPα1 sequences (BLAST 2.13.0+, with default parameters). In place of these, we added back full-length reconstructed ATPα1 sequences. RNA-seq reads were mapped to this modified transcriptome reference using bwa mem (v 0.7.17) ^57^ and processed using SAMtools (v 1.15.1) ^58^. Counts were done by htseq-count (v 2.0.2, parameters: -a 0 --nonunique all). In order to use htseq-count, a gff3 file was made for our transcriptome with the perl script gmod_fasta2gff3.pl (Key Resources Table).

We used inverted beta-binomial (ibb) tests^51^P to determine the significance of difference of expression level among tissues and paralogs, and a standard Bonferroni correction was applied to account for multiple tests (**Table S2**). To visualize the differential expression among species, tissues, and paralogs, counts were normalized using the counts function (normalized=TRUE) of the R package DeSeq2^59^. Mekko chart was plotted using ggplot2 and mekko package implemented in R.

#### CRISPR-cas9 engineered fly lines

For a list of all reagents used, see **Supplementary Table S3**. Details for each engineered fly line are as follows:

##### A119I

CRISPR-mediated mutagenesis of *D. melanogaster* was performed by WellGenetics Inc. using modified methods of Kondo and Ueda^60^: gRNA was designed and cloned into an expression vector containing a U6 promoter (pBFv-U6.2) and Cas-9 protein is supplied by a germline expression vector (pBFv-nosP-Cas9). ∼1 kilobase homology arms were amplified using Phusion High-Fidelity DNA Polymerase (Thermo Scientific) from genomic DNA of the injection strain *w^1118^*. A plasmid donor template for repair, pUC57-Kan-A119I-pBacDsRed, was constructed containing 3xP3-DsRed and two homology arms flanked by pBac terminals (“TTAA”). The substitution A119I (standardized amino acid residue numbering) was introduced into ATPα coding sequence via point mutations changing the codon “GCC” to “ATC”. Additional synonymous substitutions were added to eliminate three targeted PAM sites. For sequences of gRNA and homology arms. gRNA and donor plasmid constructs were confirmed by sequencing prior to injection. *ATP*α-targeting gRNAs and donor plasmid were microinjected into 210 embryos of the strain *w^1118^; attP40{nos-Cas9} / CyO*. Of 11 surviving G0 adults, 10 crosses to strain *w[*];;TM6B, Tb[1] / TM2, y+* were fertile and one was confirmed DsRed+. Insertion of the construct into the correct location was confirmed both by PCR and sequencing. *pBac-DsRed* was subsequently excised by first crossing to the strain *w[*]; CyO, P{Tub-pBac}/ Sp; +/TM6B* (Bloomington #8285) and subsequently to *w[*]; TM3, Sb[1] Ser[1]/TM6B, Tb[1]* (Bloomington #2537). Repeated rounds of sib-mating were used to obtain stable homozygous lines. Precise excision of *pBacDsRed* and correctly-edited genome sequence was validated by genomic PCR and sequencing.

##### Y108F-Q111E-A112S-S115L-E117T-A119N-D120N (“FESLTNN”)

Engineered line Y108F-Q111E-A112S-S115L-E117T-A119N-D120N was generated by WellGenetics Inc. (Taipei, Taiwan) using the same methods as described for A119I (including the same homology arms and gRNA). 222 *w^1118^; attP40{nos-Cas9}/CyO* embryos were injected with the *ATP*α-targeting gRNAs and donor plasmid. Of 18 surviving G0 adults, 15 were fertile in crosses to strain *w[*];;TM6B, Tb[1] / TM2, y+* and two were DsRed+. Both DsRed+ lines were PCR-sequence confirmed to contain correct insertions and processed to excise *pBac-DsRed* (as above). Each line was repeatedly sib-mated to generate homozygous lines and genome sequences were validated by genomic PCR and sequencing.

##### Q111E-A112S-A119I-D120A (“ESIA”)

Engineered line Q111E-A112S-A119I-D120A was generated by WellGenetics Inc. (Taipei, Taiwan) using the same methods as above (including homology arms and gRNA). Out of 645 microinjections of *ATP*α-targeting gRNAs, a donor plasmid and a plasmid carrying *hsp70Bb-cas9* into *w^1118^*, three lines were DsRed+ and validated carrying the correct insertion. However, after *pBac-DsRed* excision and repeated sib-mating, no homozygous mutant flies could be obtained. The bang sensitivity recovery time for ESIA/+ flies is 13.5 seconds (n=18 flies), which is comparable to flies that are heterozygous for a ATPα1 loss-of-function mutation (i.e. Δ2-6b/+, Ref. ^23^, Figure 3). Thus, this is consistent with ESIA causing loss-of-function of the *D. melanogaster* ATPα1 protein (and not mutation to a secondary site that causes lethality).

##### T797I

Engineered line T797I was generated by WellGenetics Inc. (Taipei, Taiwan) using a unique gRNA and ∼1kb homology arms. An additional mutation of a Bsal site GGTCTC → CGTCTC in the upstream homology arm in the donor plasmid was introduced to facilitate Golden Gate cloning. Of 422 injections of *ATP*α-targeting gRNAs, two lines were DsRed+ and validated as carrying the correct insertion. However, after *pBac-DsRed* excision and repeated sib-mating, no homozygous mutant flies could be obtained for either line.

##### F786Y and I787M

Engineered lines F786Y and I787M were generated in-house with CRISPR-cas9 homology-dependent repair (HDR) using one gRNA and a template single-stranded donor oligonucleotide (ssODN). The designed gRNA showed no off-target sites by *in silico* prediction (using a web-based tool called “desktop”, which is now deprecated). Oligonucleotides (IDT) were annealed to generate a T7-gRNA expression template. This template was PCR amplified, size verified on a 2% agarose gel, purified using a QIAquick spin column (Qiagen), and eluted in 30 μl Elution Buffer (Qiagen). *In vitro* transcription of gRNA templates was carried out by MEGAscript T7 Transcription Kit following the manufacturer’s protocol (Fisher Scientific). DNA and proteins were removed with turbo DNAse and phenol:chloroform:isoamyl alcohol, respectively. RNA was purified with equal volume of isopropanol, washed twice with 70% ethanol, resuspended in 30 μl RNAse-free water, and quality checked on a Bioanalyzer (Agilent). Asymmetrical ssODN design was implemented to achieve better performance ^61^. Synonymous mutations were introduced to facilitate downstream PCR screening. ssODNs were synthesized through IDT (Coralville, Iowa, USA)’s Ultramer DNA Oligo service. 20 μl mixture of 100 ng/μl gRNA and 500 ng/μl ssODNs and Cas-9 mRNA were injected into 200 embryos of the line *w^1118^; attP40{nos-Cas9} / CyO* by Rainbow Transgenic Flies, Inc (Camarillo, CA, USA). Approximately 5% of G0 offspring were fertile and were crossed to *w[*]; TM3, Sb[1] Ser[1]/TM6B, Tb[1]* (Bloomington #2537). G1 flies were separated into individual vials and again crossed to the same double balancer line (Bloomington #2537). After 3-5 days, when enough eggs were laid, the genomic DNA of G1 flies was extracted using SquishPrep protocol. A 289-bp region spanning sites 786 and 787 was PCR-amplified with primers compatible with adding customized Illumina-style i5 and i7 indexes^21^, and paired-end 150 nt sequenced on Illumina MiSeq Nano (Genomics Core Facility, Princeton, NJ, USA). Three independent lines of F786Y and two lines of I787M were obtained and confirmed by sequencing. Progeny of G1 flies with the substitution F786Y or I787M were selected and sib-mated to obtain homozygous lines.

#### Additional lines engineered using the method of Taverner et al. (2019)

Lines carrying Q111E, A112S, A119N, Q111E+A112S (“ES”) and Q111E+A112S+ A119N (“ESN”) were generated using the same method described in ref. ^23^. Substitutions, either individually or in combination, were engineered into the vector *pGX-attB-ATPατι2-6b* using Quick-change Lightning site-directed mutagenesis kit (Agilent). These plasmid constructs were injected into a white-eyed founder line *w^1118^;;ATPαΔ2-6b attP/TM6B,Tb1* by Rainbow Transgenic Flies, Inc (Camarillo, CA, USA) following their standard protocol. Lines with successfully integrated constructs (i.e. indicated by red/pink eyes) were reduced with cre-loxP excision by crossing to *y^1^, w^67c23^, P{y[+mDint2]=Crey}1b;; D*/TM3, Sb*^1^ (Bloomington #851) and succesfully reduced lines were balanced by crossing to *w*;;ry^506^ Dr^1^/TM6B, P{w[+mC]=Dfd-EYFP}3, Sb^1^,Tb^1^,ca^1^* (Bloomington 8704). Fluorescent offspring were sib-mated, and non-fluorescent *Tb^+^,Sb^+^* larvae were selected to generate stable homozygous lines. Substitutions were validated by PCR and sequencing.

#### CTS tolerance assay

Fireflies are protected by lucibufagins, a class of cardiotonic steroids (“CTS”). To measure the tolerance of engineered fly lines to CTS exposure, we introduced adult flies to food media containing varying concentrations of the representative CTS ouabain (ouabain octahydrate, Sigma-Aldrich, Cat# O-3125) and recorded how many of adults survived after 7 days. We used 0.7 grams of dried instant media (Flystuff (66-117) Nutri-fly Instant) reconstituted in a standard fly vial with 3.5 ml of 0, 5, 10, 15, or 20 mM ouabain solutions. Although the physiologically relevant concentrations of lucibufagins in fireflies are unknown, a previous study showed that wild-type flies exhibited high fatality rates upon exposure to 5 mM ouabain^23^. Reconstituted food was allowed to set for 30 minutes, and a small piece of Kimwipe tissue was added to absorb the moisture. 10 male and 10 female flies that had enclosed within 7 days were placed in each vial (three replicates per line and concentration), and kept at 25°C, 50% humidity for 7 days. Mortality rate was measured by counting the number of living flies after 7 days. Mortality was not sex-dependent, as expected based on a previous study^23^. The Cochran-Mantel-Haenzel test implemented in R was used to assess significant differences between ouabain treatments (5/10/15/20 mM) versus no ouabain.

#### Enzyme inhibition assays

CTS inhibit the ATPase activity of NKA. The principle of enzyme inhibition assays is to determine the ATPase activity of NKA by photometrically measuring the phosphate released from ATP during enzymatic hydrolysis at various concentrations of CTS. These assays were performed on wild-caught fireflies, soldier beetles and on engineered fly lines. All samples were stored at −80°C to minimize protein degradation. After thawing, fireflies and soldier beetles were immersed in deionized water and nervous tissue (brains and ventral nerve cords) was dissected under a stereomicroscope. 10 and 20 *Photinus*, 4 or 7 *Photuris*, or 10 *Rhagonycha* were pooled into each biological replicate. For flies, 90 heads were pooled together for each biological replicate (two biological replicates for *Photinus* and three for *Photuris*, *Rhagonycha*and *Drosophila*). Samples were prepared and the activities of NKA were measured following the procedures described in Taverner et al.^23^. Tissues were suspended and homogenized in deionized water using a glass grinder (Wheaton) on ice. Homogenates (split into three technical replicates for *Drosophila*), were freeze-dried (Christ, Alpha 2-4 LDPlus) overnight, and lyophilisates were reconstituted immediately before use. Samples were incubated in 6 (for beetles) or 12 (for Drosophila) increasing concentrations of ouabain (100 mM NaCl, 20 mM KCl, 4 mM MgCl_2_, 50 mM imidazol, and 2.5 mM ATP) at 37 °C for 20 minutes. A non-inhibited positive control was carried out without the addition of ouabain, and the negative control was deprived of KCI and incubated at 2 x 10^-3^ M both ouabain (inactive NKA) for complete inhibition of NKA to correct for background phosphate. Absorbance was measured at 700 nm on a CLARIOstar microplate reader (BMG Labtech, Germany). For *Drosophila*, each biological replicate was averaged over three technical replicates. Due to the limited availability of material, no technical replicates were carried out for beetle NKAs.

Relative activities of NKA were estimated for each ouabain concentration as (abs[full activity]-abs[inhibited activity])/(abs[full activity]-abs[background activity])). Curve fitting was performed with the nlsLM function from the minipack.lm library in R using the function

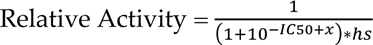

where, *x* is the ouabain concentration, *IC50* is the ouabain concentration corresponding to 50% relative activity, and *hs* is the slope coefficient. Approximate 95% confidence intervals for IC50 were estimated by parametric bootstrapping as described in ref ^23^ (**Table S4**).

For *Photuris versicolor*, the inhibition curve does not appear to be monophasic since relative enzyme activity is still ∼40% at the highest ouabain concentration. We thus made some simplifying assumptions to estimate approximate IC50s. First, it was assumed that the curve is biphasic, reflecting activities of primarily two enzyme isoforms. This can be justified by the fact that in nervous tissue (from which enzyme preps were made), paralogs ATPα1A and ATPα1C are roughly equally expressed and together account for ∼90% of total ATPα expression (**Figure 2C**). We thus used the equation

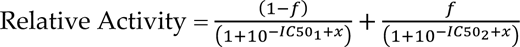

where, *x* is the ouabain concentration, *IC50_1_* and *IC50_2_* are the ouabain concentration corresponding to 50% relative activity for each isoform, respectively, *f* is the total activity attributable to the second isoform and *hs* for both enzyme isoforms is assumed to be ∼1. Approximate 95% confidence intervals for *IC50_1_* and *IC50_2_* were estimated by parametric bootstrapping as above.

#### Bang-sensitivity assay

The “bang-sensitivity” assay is a classical test where mutants with defective neurological functions experience seizures and paralysis upon mechanical over-stimulation^35,62,23,24^. Individual 14-day old male flies were placed in an empty *Drosophila* vial, vortexed at the maximum speed for 20 seconds, and immediately dumped to a surface. The time required for each individual to right itself was recorded (times >120 seconds were pooled into one timepoint). 30-50 flies were tested for each genotype. One engineered strain, ESN, was assayed in fewer individuals due to the poor condition of the flies.

### Key Resources Table

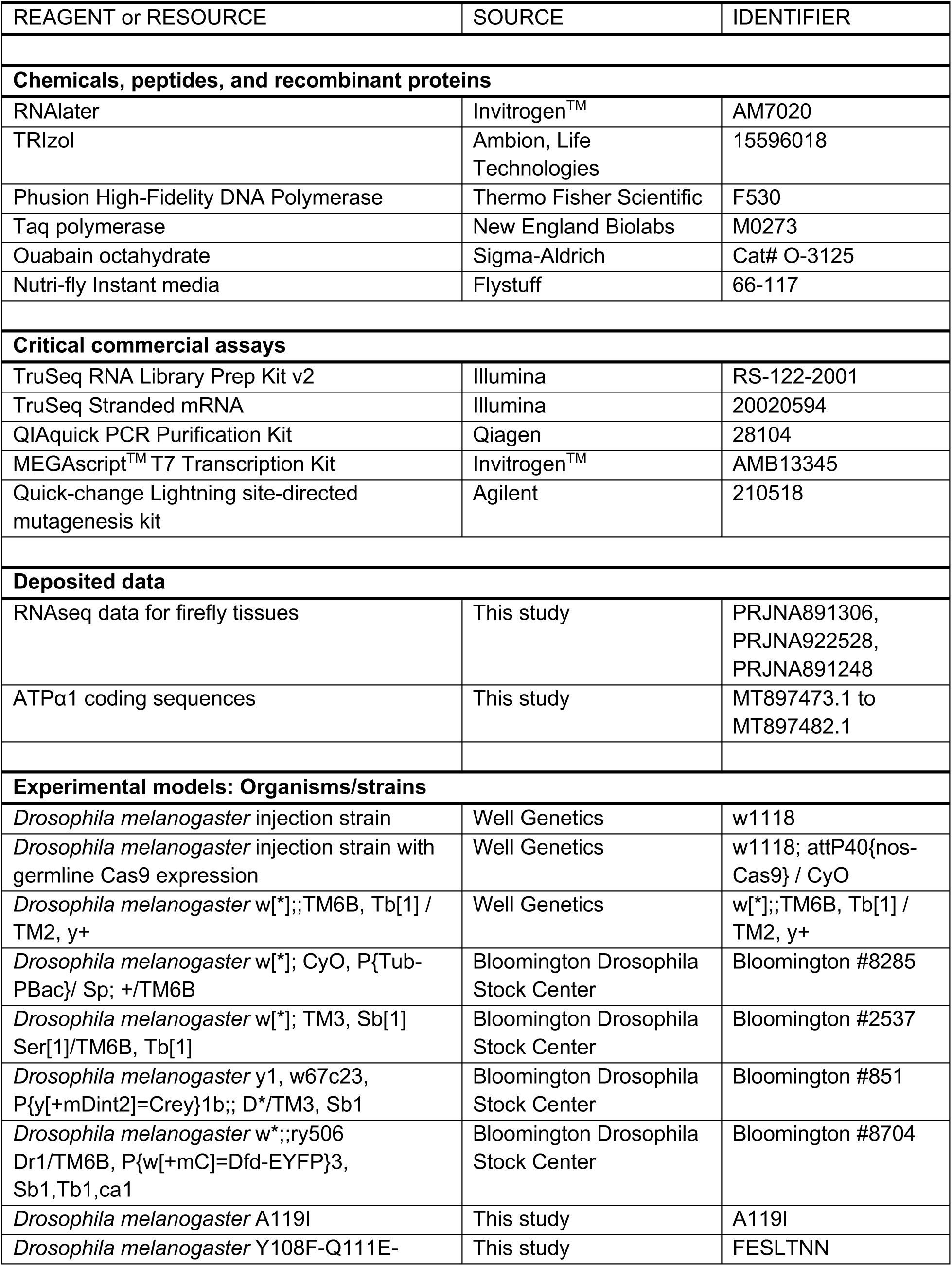
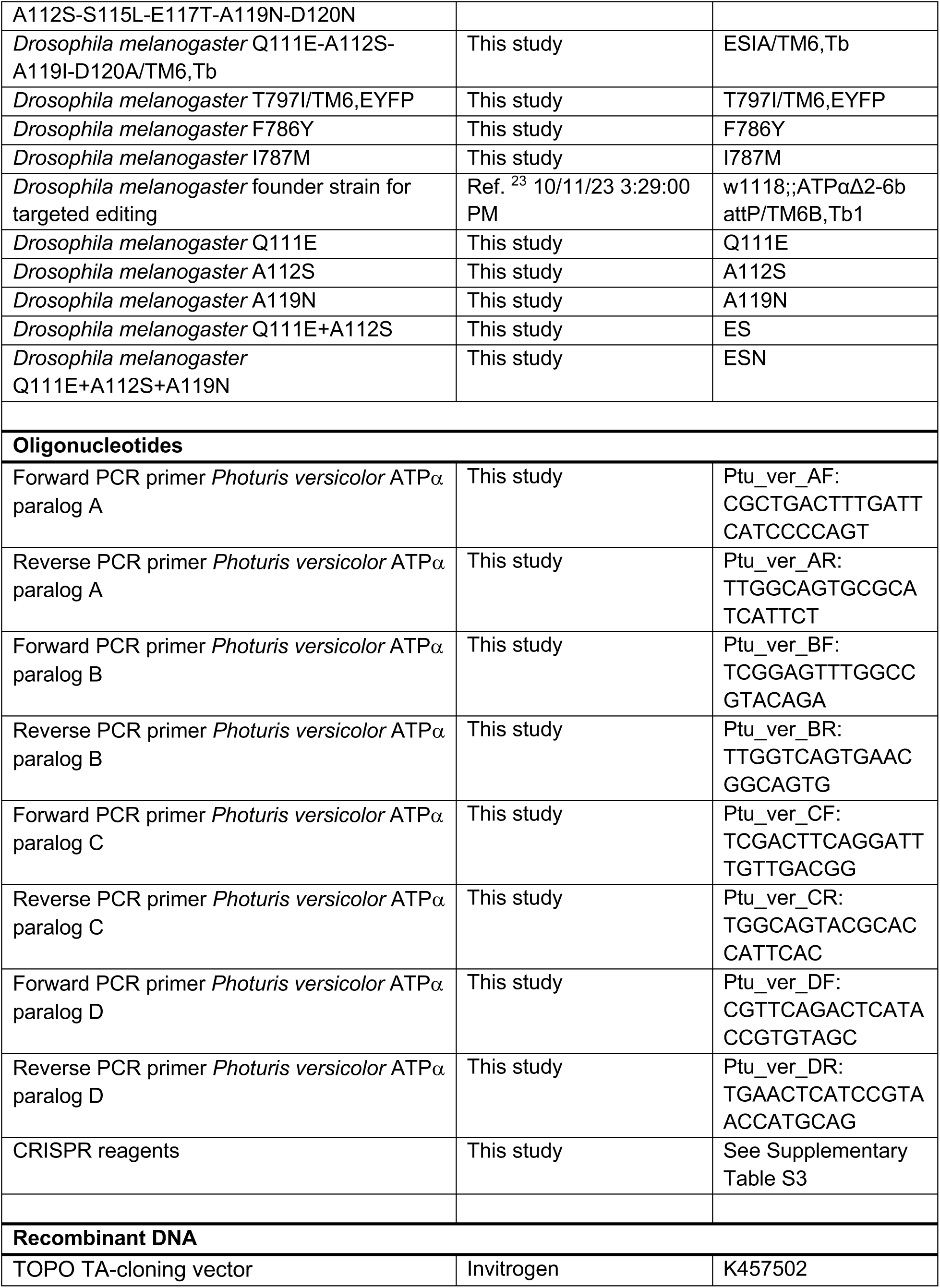
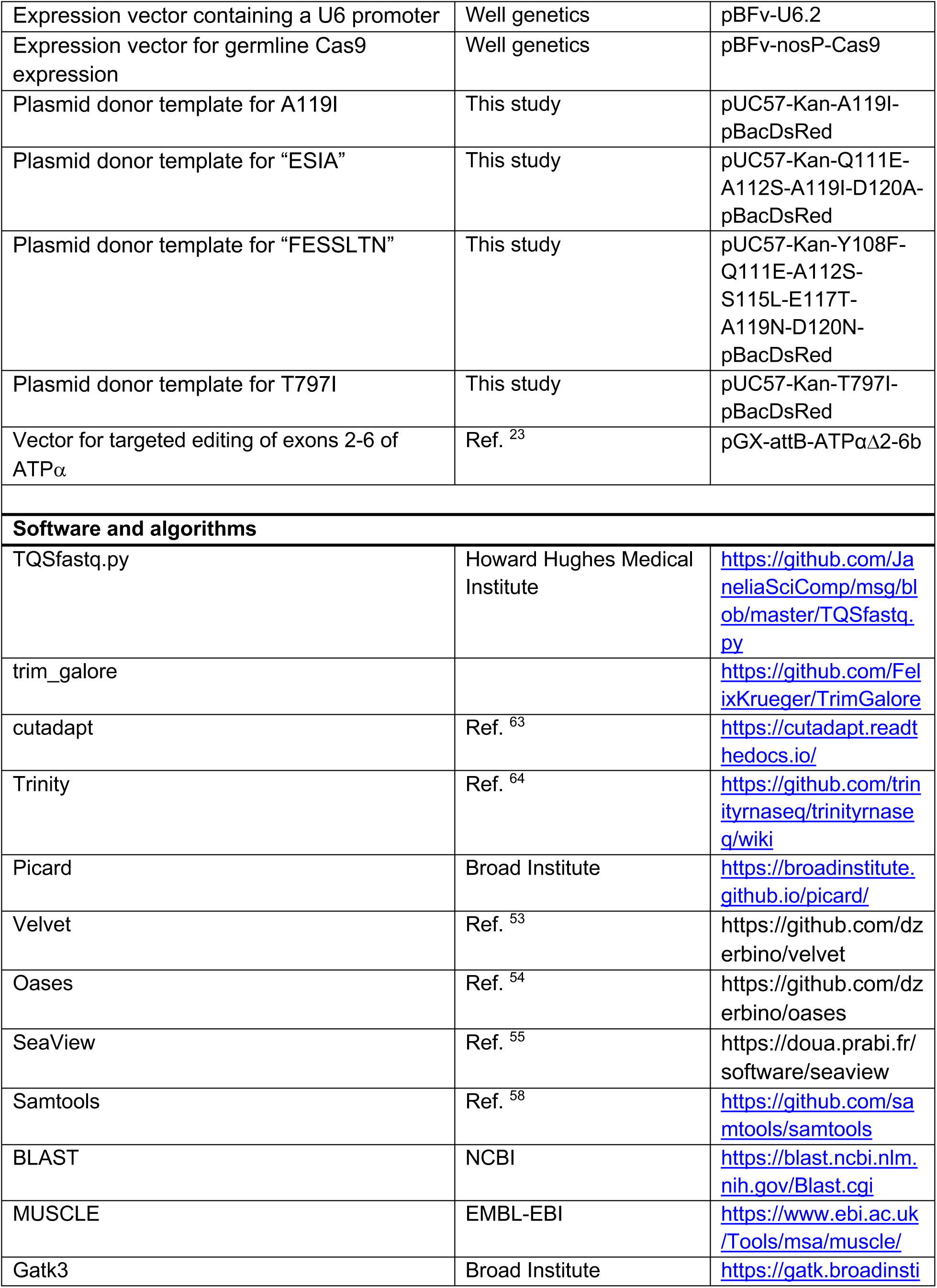
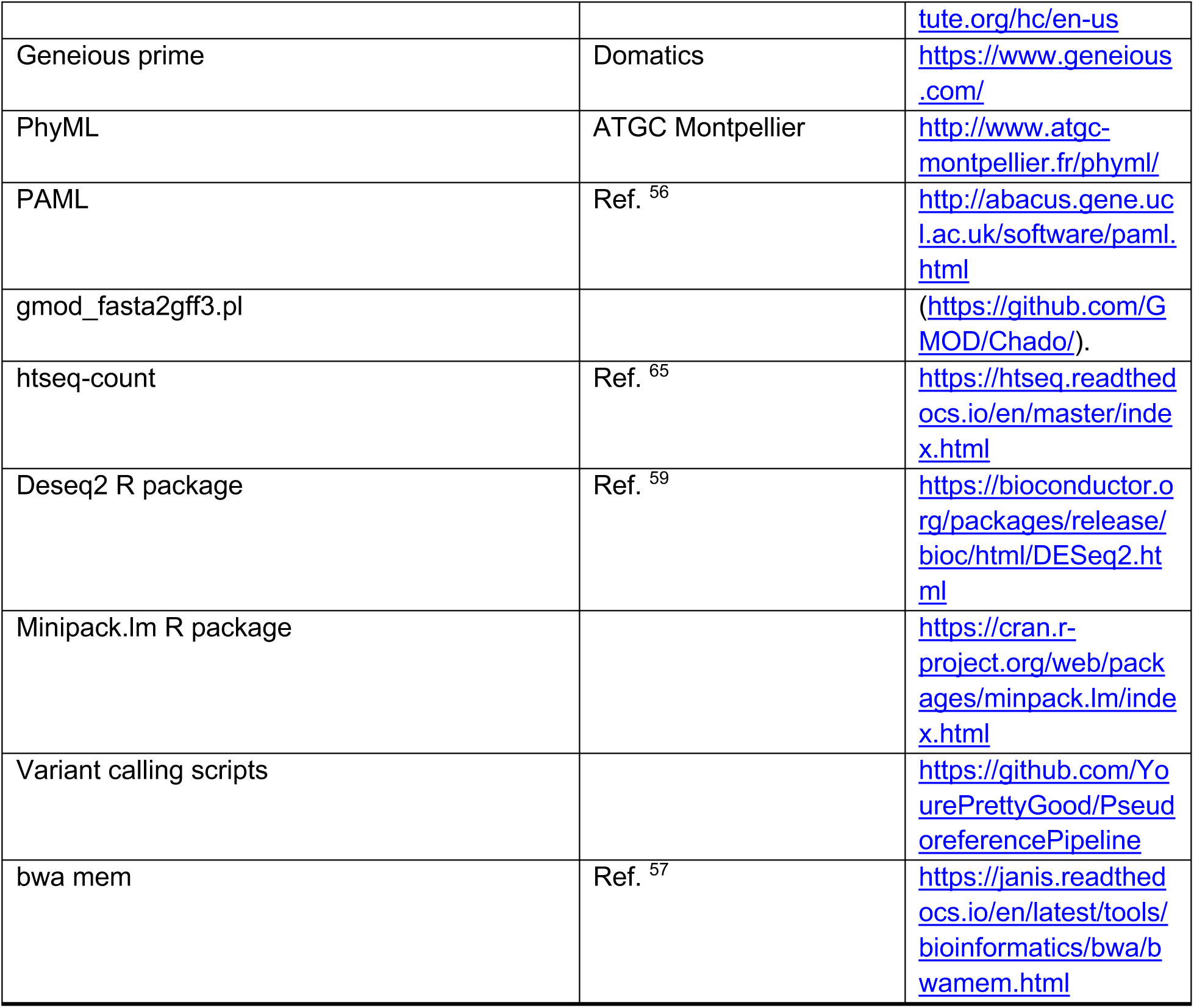

### Declaration of generative AI and AI-assisted technologies in the writing process

During the preparation of this work the author(s) used chatGPT in order to help improve the readability of some sections of this manuscript. After using this tool/service, the author(s) reviewed and edited the content as needed and take(s) full responsibility for the content of the publication.

## Supplementary Information

**Supplementary Figure S1.**
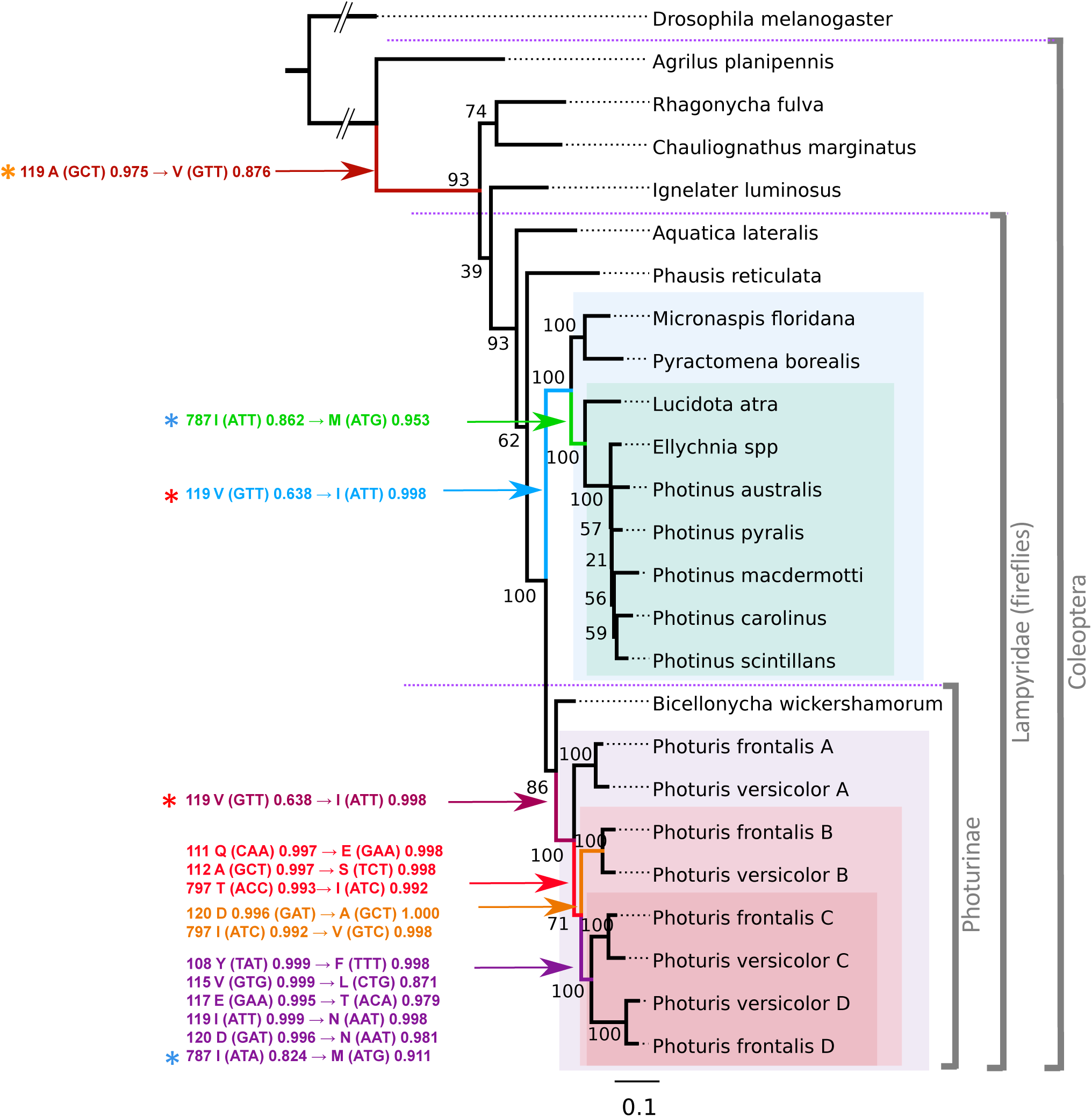
Maximum likelihood (ML) phylogeny with posterior probabilities on amino acid states (related to Figure 2). Estimation of the ML phylogeny and ancestral reconstruction implemented in PAML (Ref . S1, see Methods). The left-hand column lists the posterior probabilities of ancestral and derived states associated with key substitutions (see Methods). Only sites implicated in CTS-resistance are shown. Numbers on nodes indicate the level of bootstrap support for that particular node. A key finding is that the substitution A119V (orange asterisk) evolved prior to the diversification of predatory and prey genera whereas V119I and I787M (red and blue asterisks, respectively) evolved convergently in these genera. CTS producing firefly genera are *Lucidota*, *Ellychnia*, and *Photinus* (green shaded area).

**Supplementary Figure S2.**
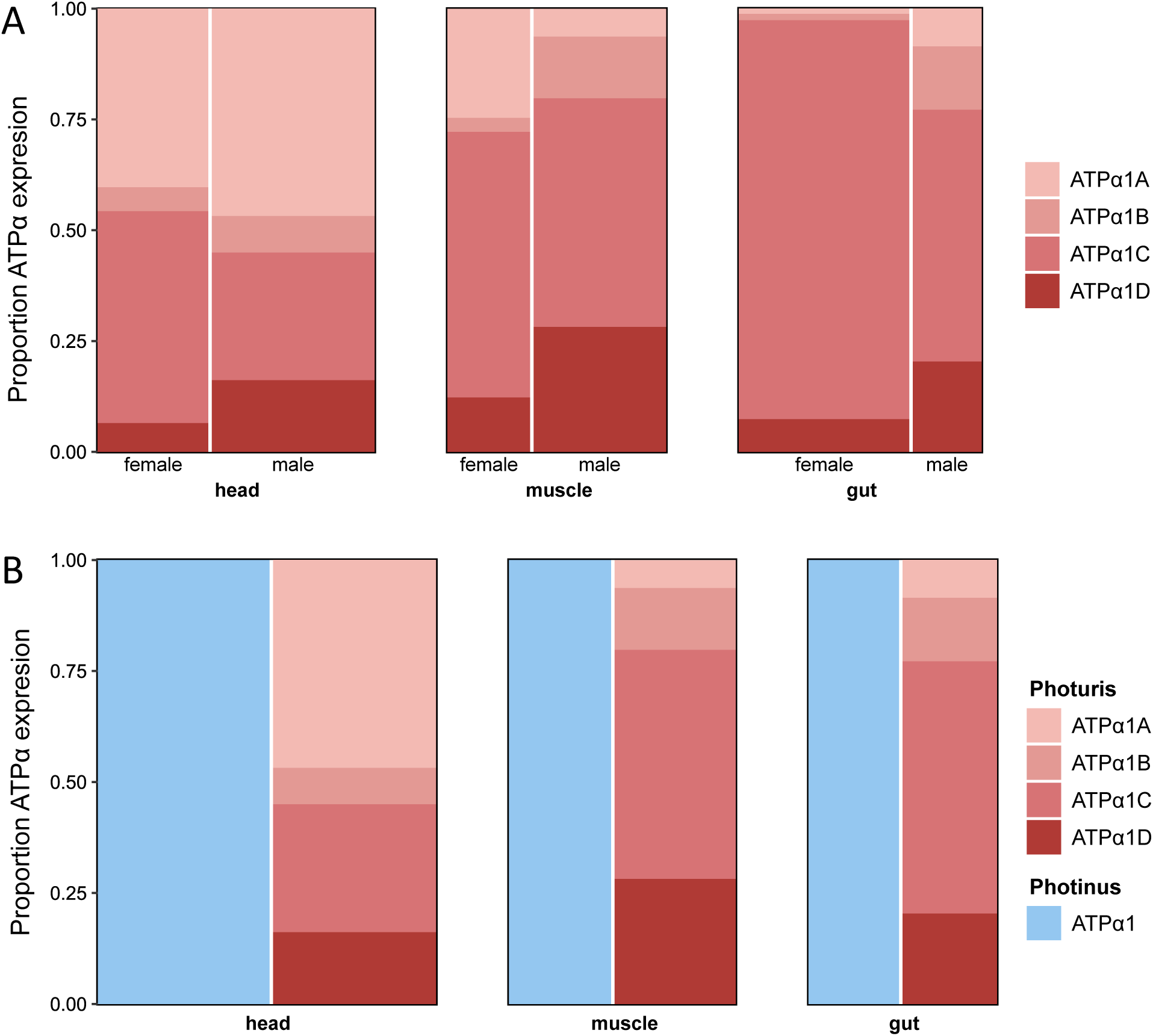
Tissue, sex and species expression patterns (related to Figure 2). (A) Tissue-specific expression of ATPα1 paralogs in *Photuris versicolor* females and males based on RNA-seq data (see Methods). Column width corresponds to relative proportion of total normalized ATPα1 expression for each tissue summed across paralogs. Shaded segments correspond to relative levels of normalized expression of the four ATPα1 paralogs within each tissue. (B) Tissue-specific expression of ATPα1 paralogs based on RNA-seq data for male *Photinus pyralis* (blue) vs *Photuris versicolor* (red) males. Column width corresponds to relative proportion of total normalized ATPα1 expression across tissues (summed across paralogs in *Photuris*). Shaded segments for *Photuris* correspond to relative levels of expression of the four ATPα1 paralogs within each tissue.

**Supplementary Figure S3.**
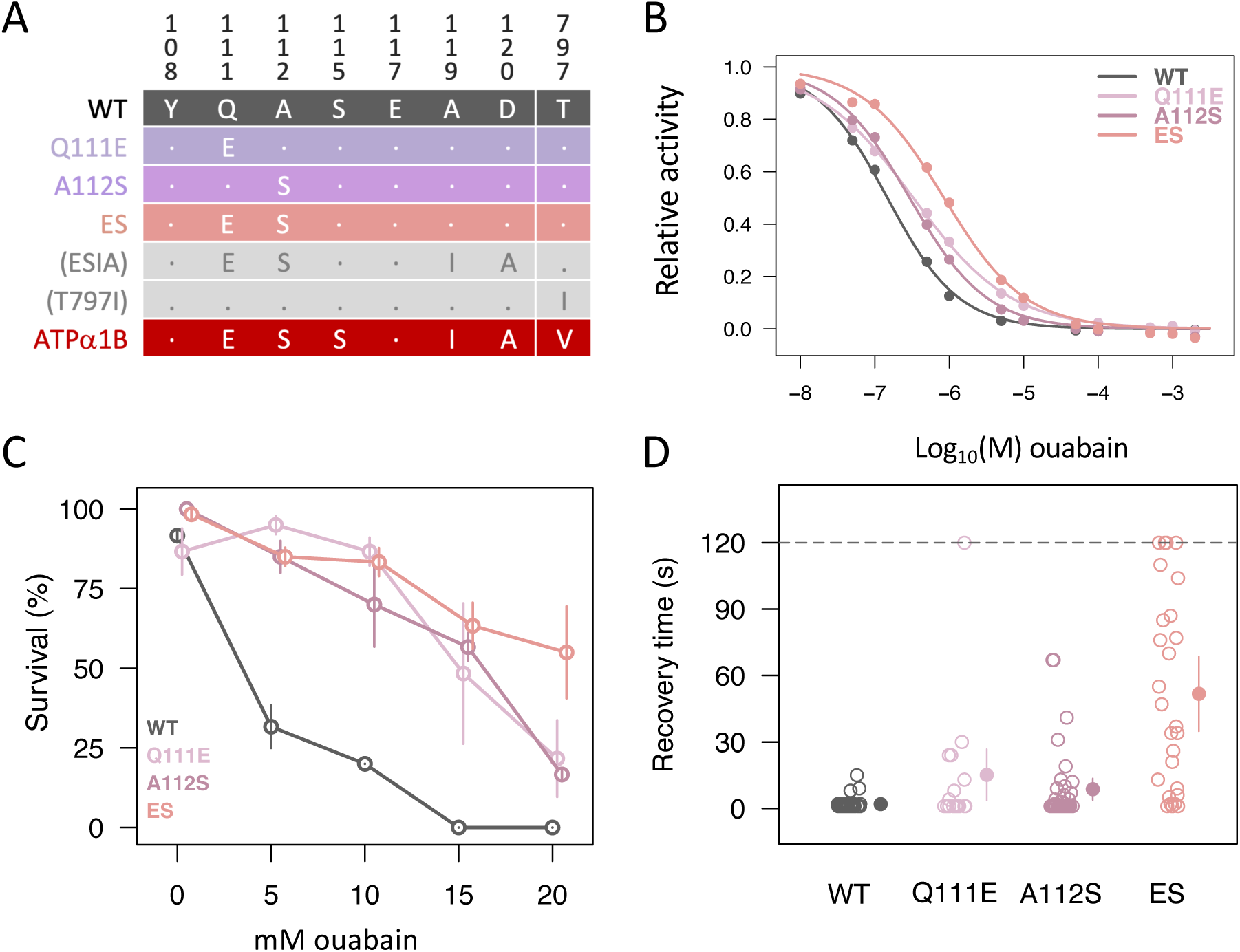
Evaluation of intermediate steps in ATPα1B evolution in *Photuris* (related to Figure 3). **(A)** Engineered fly lines carrying ATPα1B-associated substitutions, individually and in combination. Note that ESIA and T797I are homozygous lethal and were not assayed further. **(B)** CTS-inhibition assays for NKA isolated from heads of “wild type” (WT, line w^1118^) and engineered fly lines (see panel A). Mean relative activities (filled circles) are plotted as a function of increasing concentrations of the CTS ouabain. Each biological replicate (n=3, not shown) is the average of three technical replicates. Solid lines represent the least squares fit model (see **Methods**) **(C)** CTS-tolerance of adult flies. Survival of adult flies upon seven days exposure to different concentrations of CTS (0, 5, 10, 15, 20 mM ouabain). Points represent the average of three biological replicates, and bars correspond to standard errors. **(D)** Level of neural dysfunction. The assay measures recovery times following seizures induced by mechanical overstimulation (aka the “bang sensitivity” assay) of 14-day old male flies Times are capped at 120 seconds. 28-43 individuals (open circles) are assayed for each line, and the mean and 95% confidence bounds (estimated by re-sampling) are indicated as points and whiskers, respectively.

**Supplementary Figure S4.**
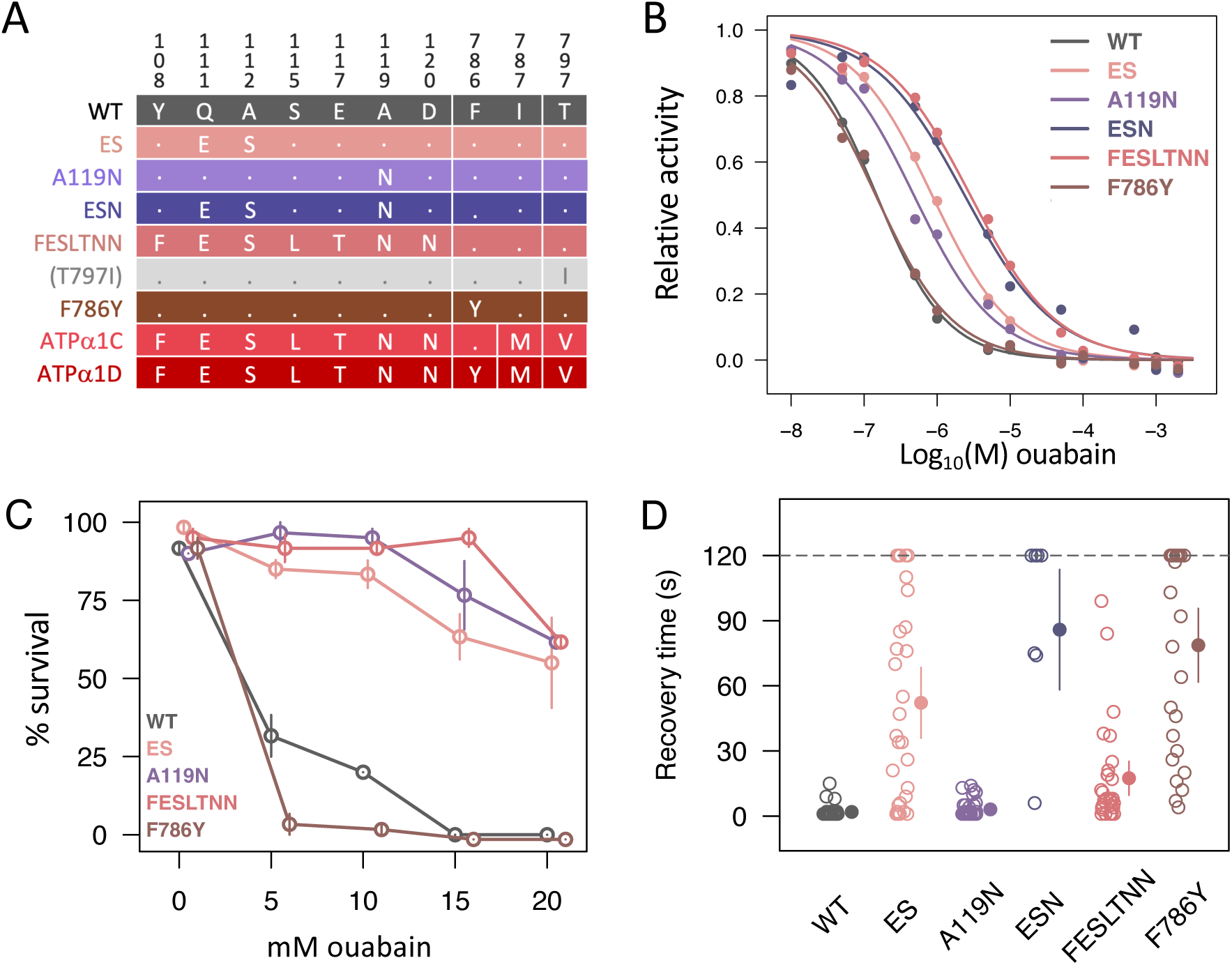
Evaluation of amino acid substitutions contributing to CTS resistance of ATPα1 paralogs C and D in *Photuris* (related to Figure 3). **(A)** A collection of engineered fly lines carrying substitutions occurring along the ATPα1C and D lineages. The color code follows that in the main text. T797I was homozygous lethal and not assayed further. Results for I787M can be found in Figure 4. **(B)** CTS-inhibition assays for NKA isolated from heads of w^1118^ (WT) and engineered fly lines. Mean relative activities (filled circles) are plotted as a function of increasing concentrations of the CTS ouabain. Each biological replicate (n=3, not shown) is the average of three technical replicates. Solid lines represent the least squares fit model (see **Methods**) **(C)** CTS-tolerance of adult flies. Survival of adult flies upon seven days exposure to different concentrations of CTS (0, 5, 10, 15, 20 mM ouabain). Points represent the average of three biological replicates, and bars correspond to standard errors. **(D)** Level of neural dysfunction. The assay measures recovery times following seizures induced by mechanical overstimulation (aka “bang sensitivity” assay) of 14-day old male flies (times capped at 120 seconds). 28-43 individuals (open circles) are assayed for each line, and the mean and 95% confidence bounds (estimated by re-sampling) are indicated as points and whiskers, respectively.

**Supplementary Figure S5.**
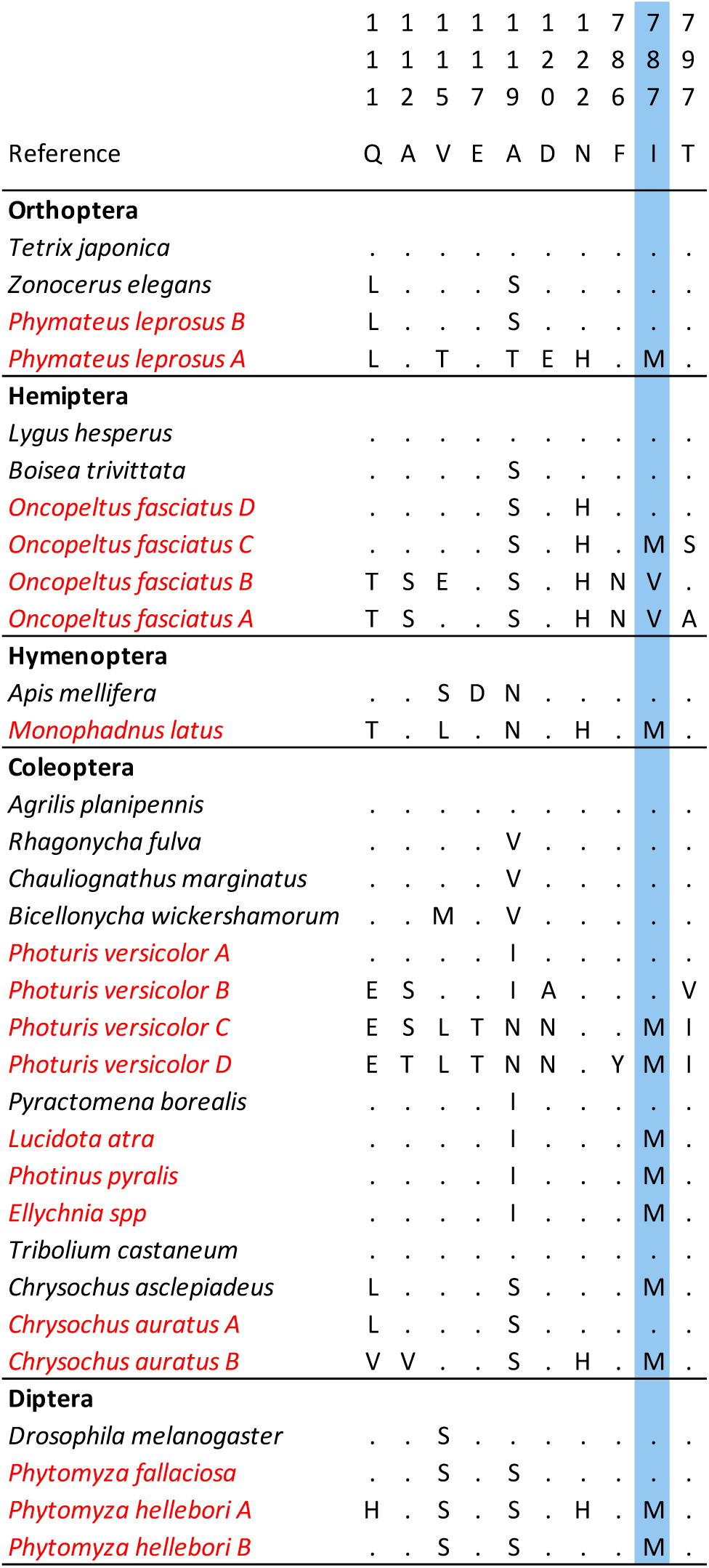
Amino acid states at key sites in ATPα1 of CTS-associated insects and representative outgroups (related to Figure 4). Amino acid numbering follows the standard numbering scheme based on the sheep (*Ovis ares*) ATP1A1 sequence (see Ref S2). CTS-associated insects are highlighted in red. The substitution I787M (at highlighted site 787) appears repeatedly in CTS-associated taxa.

**Table S1.**
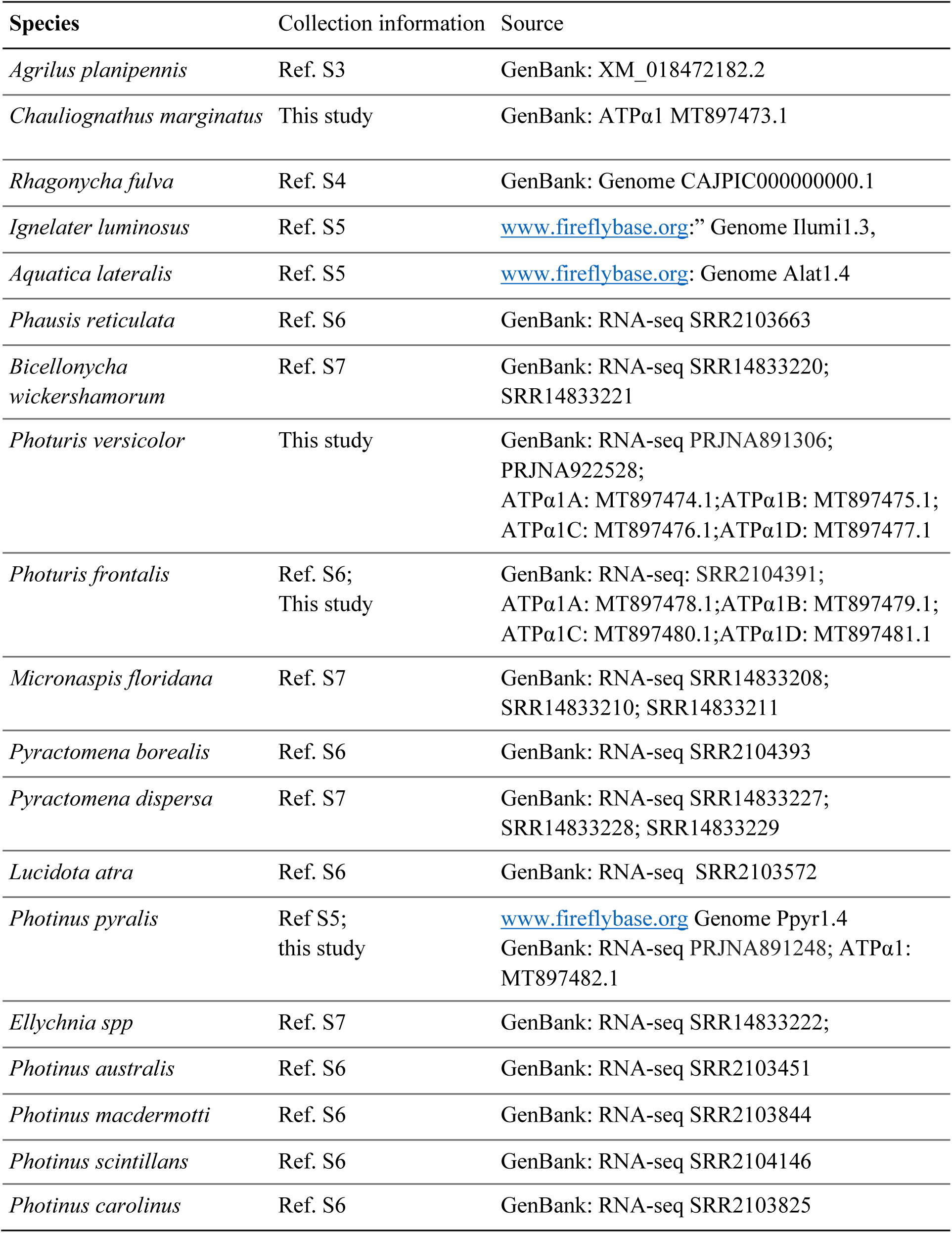
Sources of sequence data used in this study.

**Table S2.**
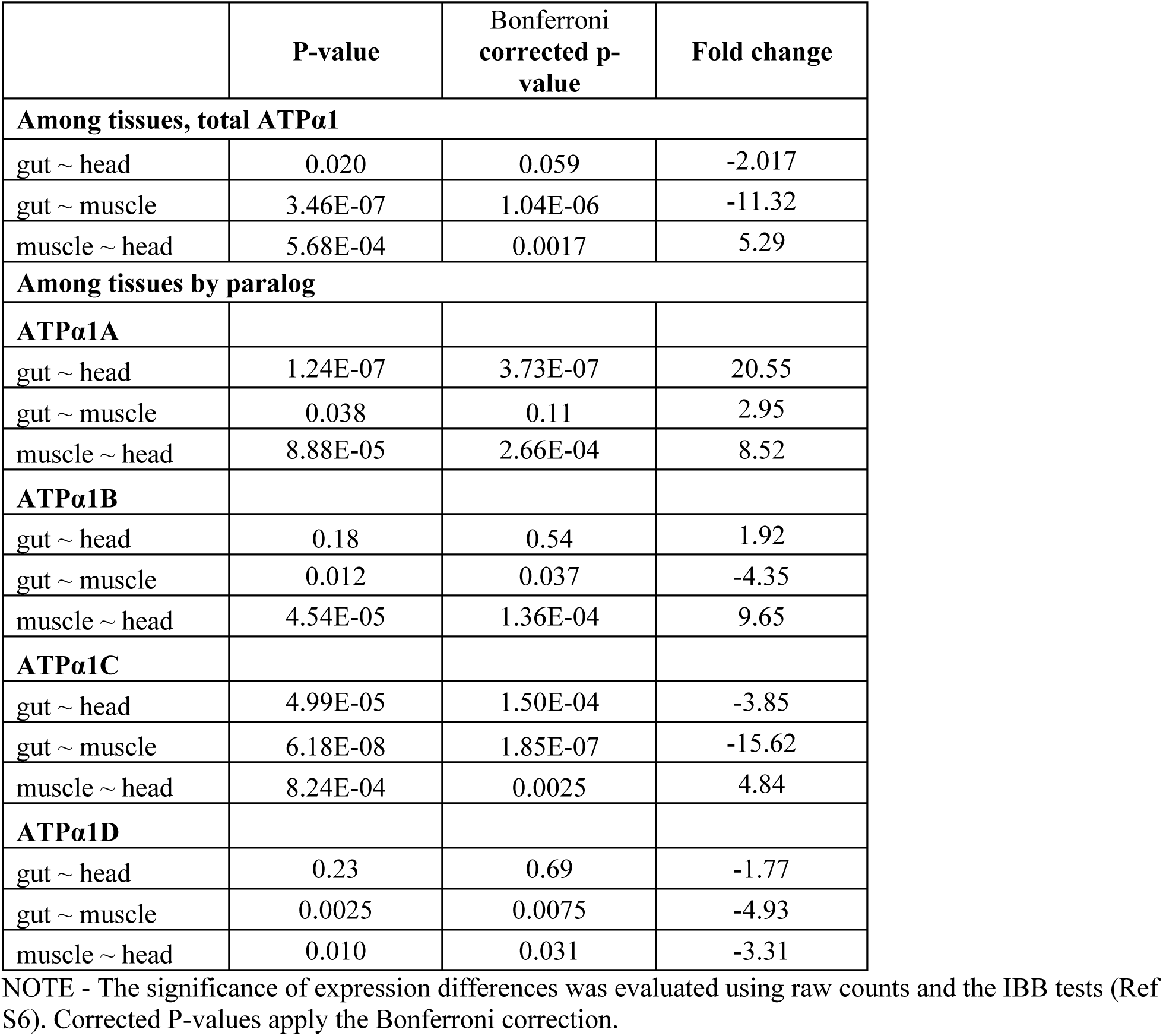
Differential expression of ATPα1 paralogs among tissues in *Photuris*.

|  | **P-value** | **Bonferroni**<br>**corrected p-**<br>**value** | **Fold change** |
| --- | --- | --- | --- |
| **Among tissues, total ATPα1** |  |  |  |
| gut ~ head | 0.020 | 0.059 | -2.017 |
| gut ~ muscle | 3.46E-07 | 1.04E-06 | -11.32 |
| muscle ~ head | 5.68E-04 | 0.0017 | 5.29 |
| **Among tissues by paralog** |  |  |  |
| **ATPα1A** |  |  |  |
| gut ~ head | 1.24E-07 | 3.73E-07 | 20.55 |
| gut ~ muscle | 0.038 | 0.11 | 2.95 |
| muscle ~ head | 8.88E-05 | 2.66E-04 | 8.52 |
| **ATPα1B** |  |  |  |
| gut ~ head | 0.18 | 0.54 | 1.92 |
| gut ~ muscle | 0.012 | 0.037 | -4.35 |
| muscle ~ head | 4.54E-05 | 1.36E-04 | 9.65 |
| **ATPα1C** |  |  |  |
| gut ~ head | 4.99E-05 | 1.50E-04 | -3.85 |
| gut ~ muscle | 6.18E-08 | 1.85E-07 | -15.62 |
| muscle ~ head | 8.24E-04 | 0.0025 | 4.84 |
| **ATPα1D** |  |  |  |
| gut ~ head | 0.23 | 0.69 | -1.77 |
| gut ~ muscle | 0.0025 | 0.0075 | -4.93 |
| muscle ~ head | 0.010 | 0.031 | -3.31 |

NOTE - The significance of expression differences was evaluated using raw counts and the IBB tests (Ref S6). Corrected P-values apply the Bonferroni correction.

**Table S3.**
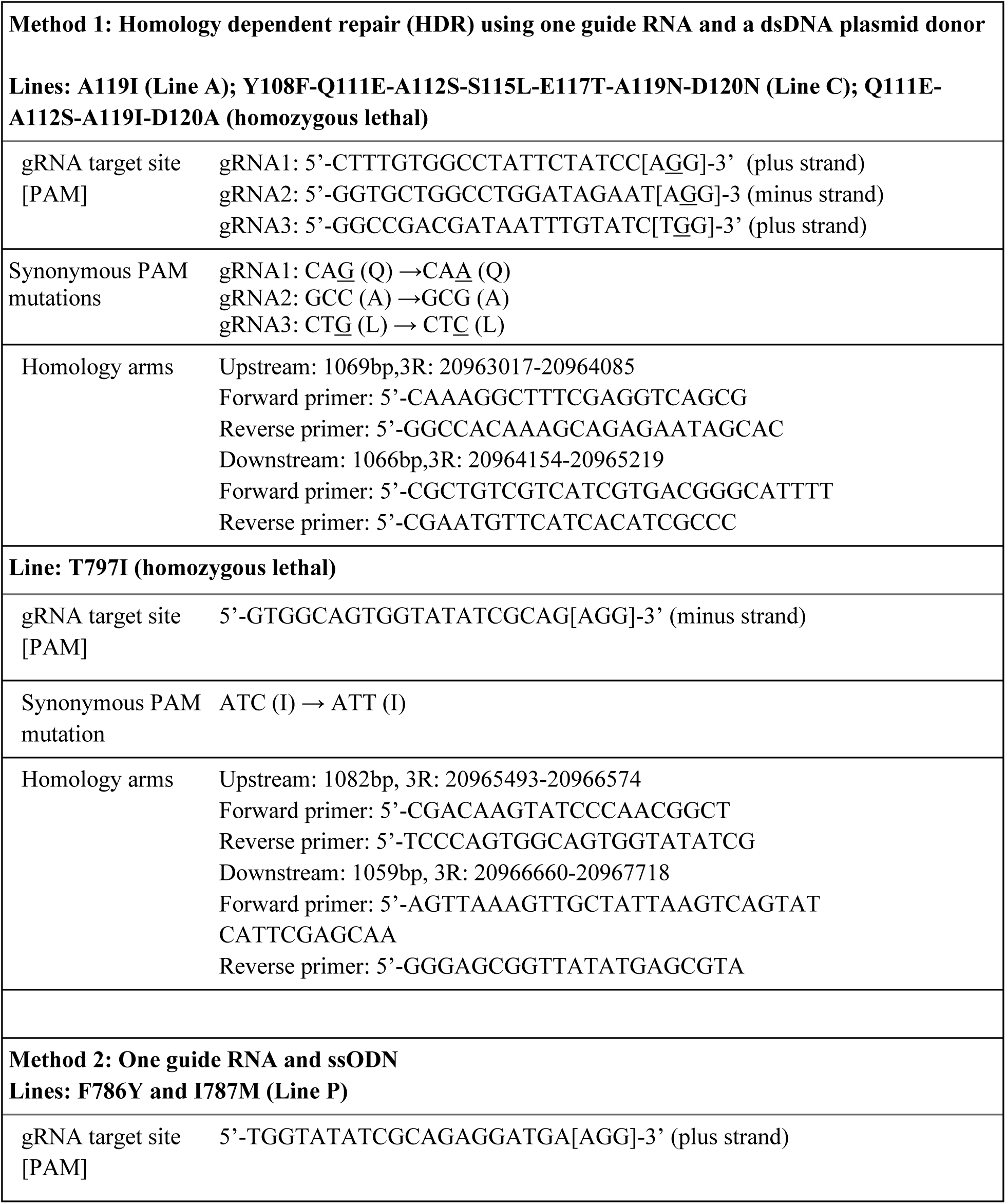
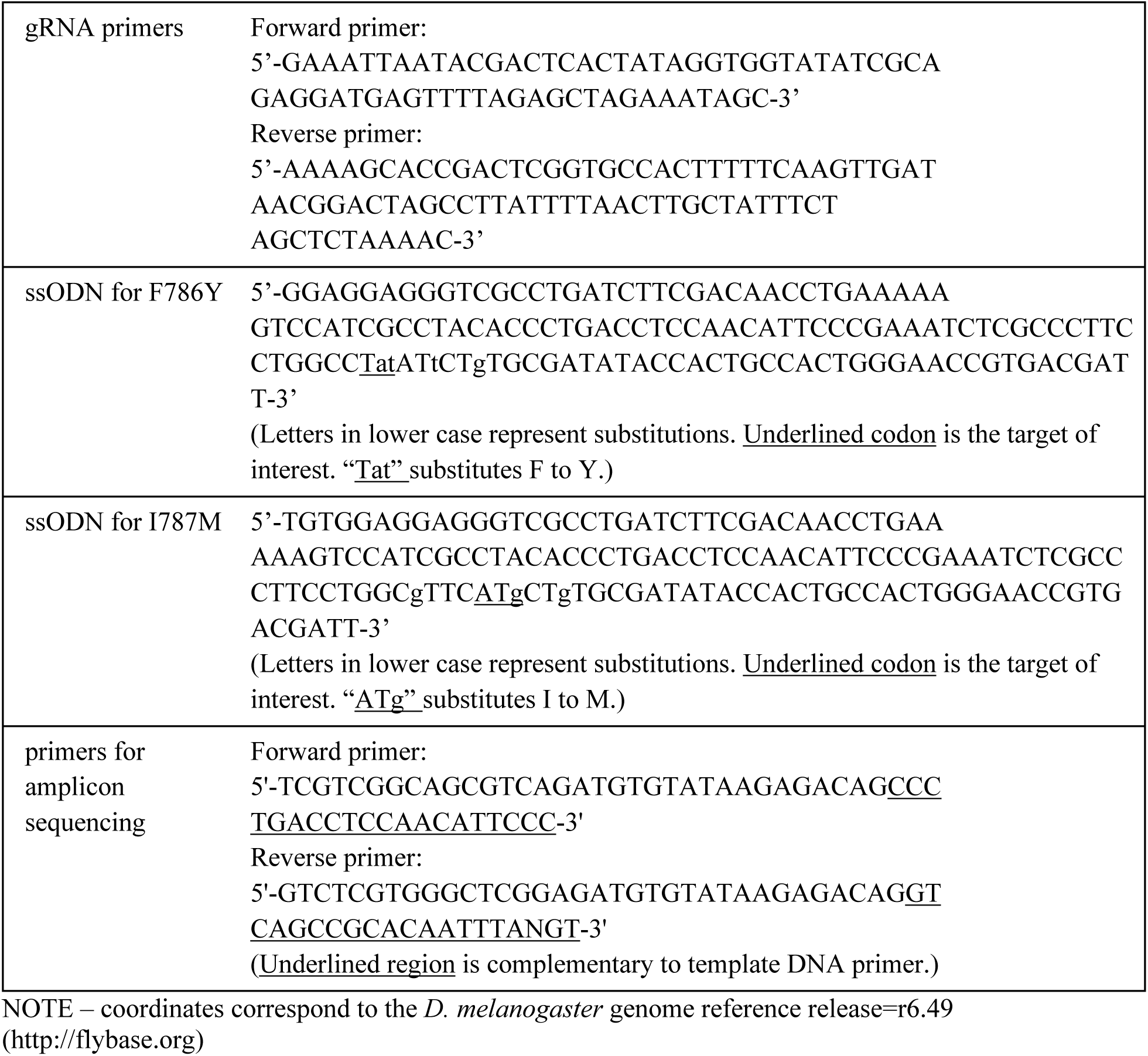
Reagents used for CRISPR-mediated mutagenesis of *Drosophila* ATPa.

**Table S4.**
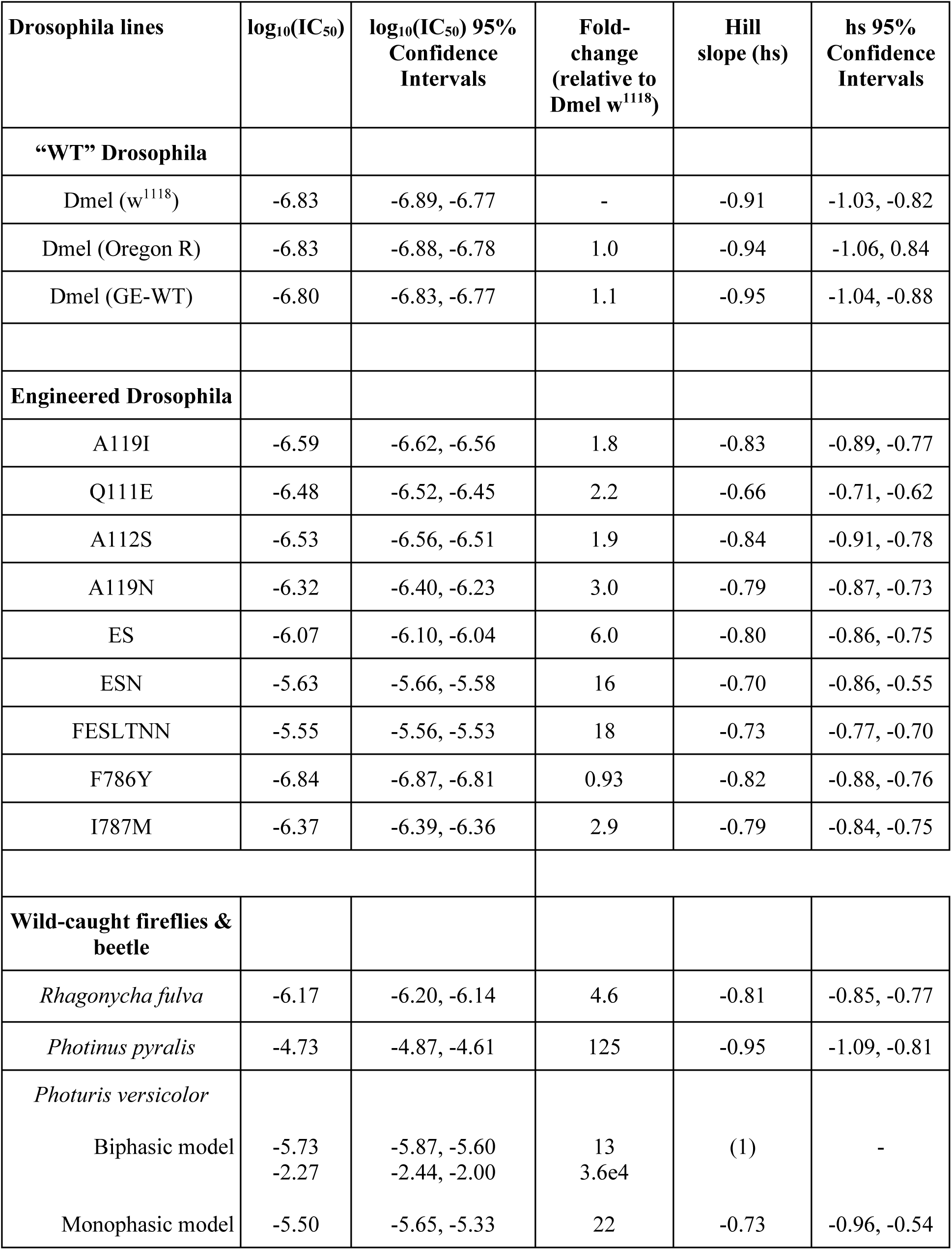
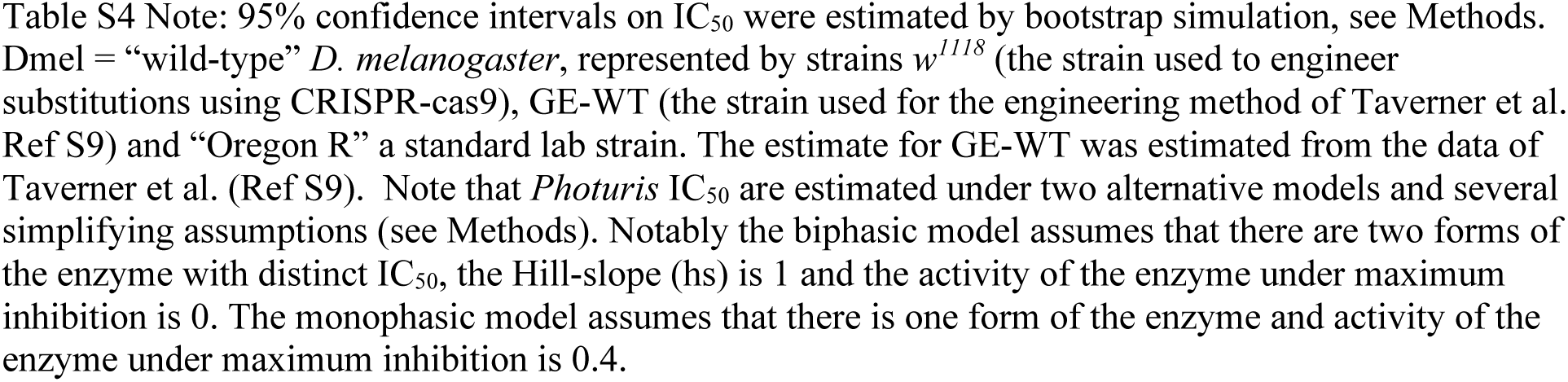
Estimates of CTS-resistance of Na^+^,K^+^-ATPase (IC50 of ouabain).

| **Drosophila lines** | **log<sub>10</sub>(IC<sub>50</sub>)** | **log<sub>10</sub>(IC<sub>50</sub>) 95% Confidence Intervals** | **Fold-change (relative to Dmel w<sup>1118</sup>)** | **Hill slope (hs)** | **hs 95% Confidence Intervals** |
| --- | --- | --- | --- | --- | --- |
| **“WT” Drosophila** |  |  |  |  |  |
| Dmel (w<sup>1118</sup>) | -6.83 | -6.89, -6.77 | - | -0.91 | -1.03, -0.82 |
| Dmel (Oregon R) | -6.83 | -6.88, -6.78 | 1.0 | -0.94 | -1.06, 0.84 |
| Dmel (GE-WT) | -6.80 | -6.83, -6.77 | 1.1 | -0.95 | -1.04, -0.88 |
| **Engineered Drosophila** |  |  |  |  |  |
| A119I | -6.59 | -6.62, -6.56 | 1.8 | -0.83 | -0.89, -0.77 |
| Q111E | -6.48 | -6.52, -6.45 | 2.2 | -0.66 | -0.71, -0.62 |
| A112S | -6.53 | -6.56, -6.51 | 1.9 | -0.84 | -0.91, -0.78 |
| A119N | -6.32 | -6.40, -6.23 | 3.0 | -0.79 | -0.87, -0.73 |
| ES | -6.07 | -6.10, -6.04 | 6.0 | -0.80 | -0.86, -0.75 |
| ESN | -5.63 | -5.66, -5.58 | 16 | -0.70 | -0.86, -0.55 |
| FESLTNN | -5.55 | -5.56, -5.53 | 18 | -0.73 | -0.77, -0.70 |
| F786Y | -6.84 | -6.87, -6.81 | 0.93 | -0.82 | -0.88, -0.76 |
| I787M | -6.37 | -6.39, -6.36 | 2.9 | -0.79 | -0.84, -0.75 |
| **Wild-caught fireflies & beetle** |  |  |  |  |  |
| *Rhagonycha fulva* | -6.17 | -6.20, -6.14 | 4.6 | -0.81 | -0.85, -0.77 |
| *Photinus pyralis* | -4.73 | -4.87, -4.61 | 125 | -0.95 | -1.09, -0.81 |
| *Photuris versicolor* |  |  |  |  |  |
| Biphasic model | -5.73<br>-2.27 | -5.87, -5.60<br>-2.44, -2.00 | 13<br>3.6e4 | (1) | - |
| Monophasic model | -5.50 | -5.65, -5.33 | 22 | -0.73 | -0.96, -0.54 |

